# SARS-CoV-2 NSP14 inhibitor exhibits potent antiviral activity and reverses NSP14-driven host modulation

**DOI:** 10.1101/2025.08.21.671674

**Authors:** Mengxin Luo, Jun Mo, Ziqiao Wang, Huimin Wei, Kexin Chen, Liteng Shen, Ying Wang, Linjie Li, Yongkang Chen, Weihao Chen, Xue Li, Hui Feng, Xinyu Wang, Huan Zhou, Bizhi Li, Feng Xu, Qingwei Zhao, Yichen Xu, Jinxin Che, Peng Zou, Rong Zhang, Xiaowu Dong, Wei Xie

## Abstract

The emergence of SARS-CoV-2 variants and drug-resistant mutants highlights the urgent need for novel antiviral therapeutics. SARS-CoV-2 NSP14, an N7-guanosine methyltransferase, plays a critical role in viral RNA capping, enabling viral replication and immune evasion. While NSP14 has emerged as a promising drug target, its role in host-virus crosstalk and the cellular consequences of NSP14 inhibition remain poorly understood. Here, we present the identification and characterization of C10, a highly potent and selective first-in-class non-nucleoside inhibitor of the NSP14 S-adenosylmethionine (SAM)-binding pocket. C10 demonstrates robust antiviral activity against SARS-CoV-2, including its variants, with EC_50_ values ranging from 64.03 to 301.9 nM, comparable to the FDA-approved drug remdesivir in our cell-based assays. C10 also exhibits broad-spectrum activity against other betacoronaviruses and inhibits SARS-CoV-2 at the replication stage. C10 suppresses viral translation and exhibits immunostimulatory effect. Additionally, C10 specifically reversed NSP14-mediated alterations in host transcriptome. The antiviral efficacy of C10 was further validated in a transgenic mouse model of SARS-CoV-2 infection. Our findings highlight C10 as a promising candidate for the development of effective treatments against SARS-CoV-2 and its emerging variants. This study also uncovers a novel mechanism of NSP14 in SARS-CoV-2 pathogenesis and its therapeutic potential, providing insights that may extend to other viral capping methyltransferases.

## Main

The COVID-19 pandemic has spurred an urgent pursuit of effective antiviral treatments^1,2^. FDA-approved drugs directly targeting SARS-CoV-2 focus on inhibiting the viral main protease (Mpro) or RNA-dependent RNA polymerase^3^; however, drug-resistant SARS-CoV-2 strains have been identified in COVID-19 patients^4,5^. Therefore, additional antivirals with alternative mechanisms of action are urgently required to combat drug-resistant and emerging SARS-CoV-2 variants.

Coronaviruses employ their own RNA capping machinery to synthesize the 5′-end cap of viral RNAs in the cytoplasm^6^, mimicking host mRNA to promote viral replication and evade innate immune sensing^7–10^. SARS-CoV-2 nonstructural protein 14 (NSP14) is a bifunctional enzyme composed of two distinct domains: a 3′–5′ exoribonuclease (ExoN) domain for viral RNA proofreading^11,12^ and an N7-guanosine methyltransferase (N7-MTase) domain that catalyzes the methylation of the guanine-N7 position of viral RNA^13^. Notably, mutagenesis and structural studies suggest a strong functional interdependence between these two domains, by forming a tightly coupled structural framework that supports coordinated regulation of viral RNA proofreading and capping^14–16^. Loss-of-function mutations in either domain have been shown to severely impair viral viability^17–19^, highlighting the essential role of NSP14 in viral replication and its potential as a therapeutic target^11,20–22^. In addition, the individual expression of NSP14 drives host transcriptome modulation to favor SARS-CoV-2 infection^23^, implying the pathogenic role of NSP14 beyond viral RNA capping.

The catalytic pocket of NSP14 N7-MTase includes the S-adenosylmethionine (SAM, the universal methyl donor)-binding site and the N7-guanosine RNA cap-binding site. The structural features of NSP14 within its SAM-binding pocket are distinct from those of human methyltransferases, make it an ideal target for highly selective drug design^14,24–29^. Several nucleotide-based inhibitors of NSP14’s SAM-binding pocket have been reported, including adenosine mimetics and bisubstrate analogs^30–38^. While these compounds demonstrate potent *in vitro* inhibition of N7-MTase activity, their efficacy in infected cells remains limited. In contrast, the reported non-nucleotide NSP14 inhibitors have generally exhibited lower biopotency and limited target selectivity^39–43^. Until very recently, a first-in-class small molecule inhibitor targeting the NSP14 RNA cap-binding site was disclosed, demonstrating promising antiviral activity and significant therapeutic potential^44^. However, highly potent non-nucleotide inhibitors specifically targeting the SAM-binding pocket of NSP14 with *in vivo* efficacy have not yet been reported. Moreover, the pharmacological effects of NSP14 inhibitors on host-virus interactions and their underlying mechanisms are still poorly understood.

In this study, we discovered and developed non-nucleotide SARS-CoV-2 NSP14 inhibitors that specifically occupy the SAM-binding site. We uncovered the multiple antiviral effects of NSP14 inhibition, including suppression of viral translation, immunostimulatory effect, and reversing host transcriptome modulation. The antiviral efficacy of the lead compounds was validated in both a cell-based assays and SARS-CoV-2-infected mice. Our data indicate that NSP14 inhibition holds promise for developing effective treatments against SARS-CoV-2 and emerging variants.

## Results

### Discovery of NSP14 inhibitors by structure-guided virtual screening

To identify SARS-CoV-2 NSP14 inhibitors, we established a structure-guided virtual screening against NSP14 N7-MTase activity. Approximately 150,000 compounds were docked to the SAM-binding site of the NSP14 structure (PDB ID 7R2V) using high-throughput virtual screening (**Fig. 1a**). The compounds were refined to form an intermediate library of 200 top-ranked compounds using docking simulations that involved the application of standard precision and extra precision modes. The DeepDock program, a deep learning framework for protein-ligand docking^45^, was then applied for further screening (**Fig. 1a**). After filtration based on drug-likeness prediction and clustering analysis, 20 compounds were selected for further NSP14 inhibition evaluation.

**Figure 1.**
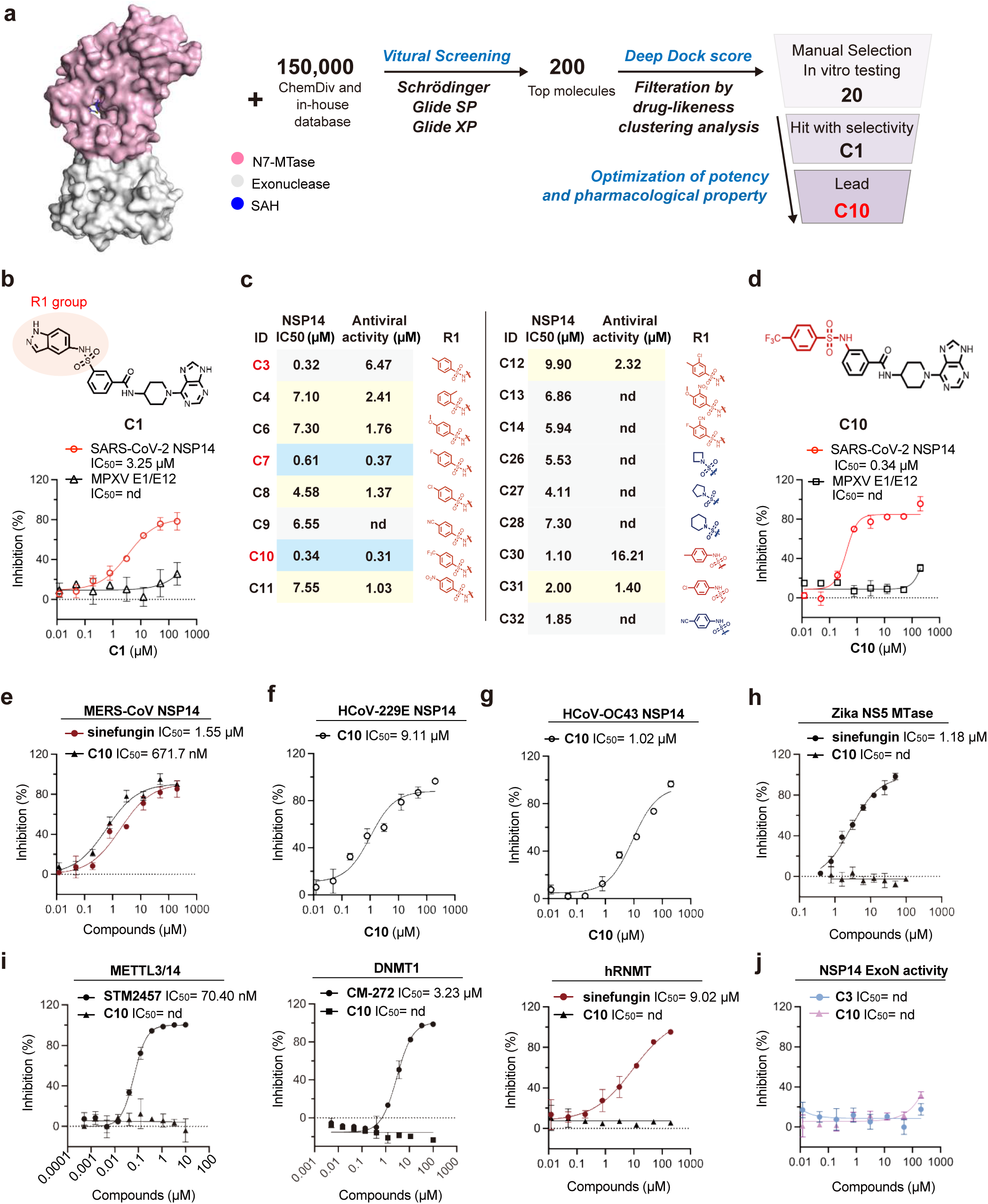
Discovery and Characterization of Compound C10 Targeting NSP14 SAM-Binding Pocket. **a.** Flowchart of the virtual screening and development of C chemotype inhibitors. **b.** Chemical structure of hit compound C1 (Left). Dose-dependent inhibition curves of C1 against SARS-CoV-2 NSP14 and MPXV E1/E12 as measured by the MTase-Glo assay (Right). Data are presented as mean ± SD from three independent replicates. **c.** Bioactivity chart of selected C chemotype inhibitors in the MTase-Glo assay and antiviral assessment using SARS-CoV-2. **d.** Chemical structure of hit compound C10 (Left). Dose-dependent inhibition curves of C1 against SARS-CoV-2 NSP14 and MPXV E1/E12 as measured by the MTase-Glo assay (Right). Data are presented as mean ± SD from three independent replicates. **e-g.** Dose-response inhibition curves of C10 and reference compound sinefungin against MERS-CoV NSP14 (e), HCoV-229E NSP14 (f), and HCoV-OC43 NSP14 (g) as measured by the MTase-Glo assay. Data are mean ± SD from three independent replicates. **h.** Dose-response inhibition curves of C10 and sinefungin against Zika virus NS5 methyltransferase, measured by the FP assay. Data are mean ± SD from three independent replicates. **i.** Selectivity profile of C10 against human methyltransferases: METTL3/14 (compared with STM2457), DNMT1 (compared with CM-272), and RNMT (compared with sinefungin). Data are mean ± SD from three independent replicates. **j.** Dose-response inhibition curves of C3 and C10 against the NSP14-NSP10 complex, measured by the FRET-based exonuclease assay. Data are mean ± SD from three independent replicates.

The dose-response evaluations of these compounds for inhibiting NSP14 methyltransferase activity were conducted using a luminescence-based enzymatic assay, which utilized 300 nM NSP14, 1.5 μM GpppG cap, and 1.5 μM methyl donor S-adenosylmethionine (SAM). Compound C1 showed the strongest inhibitory activity against NSP14, with a half-maximal inhibitory concentration (IC_50_) value of 3.25 µM (**Fig. 1b**). The target specificity of C1 was primarily validated as it showed minor inhibitory effects on monkeypox virus (MPXV) E1/E12 N7-MTase in the counter-screen, establishing it as a promising hit compound (**Fig. 1b**). After optimization of potency in enzymatic assay and SARS-CoV-2 infection (**Fig. 1c, Extended Data Fig. 1**), C3 was selected by showing a 10-fold improvement in IC_50_ value (0.32 µM) and enhanced target specificity (**Extended Data Fig. 2a-c**). To enhance the pharmacological properties of C3, we developed the lead compound C10 (IC_50_ = 0.34 µM) for further characterization and functional testing (**Fig. 1d**). To further evaluate C10’s inhibitory effect on N7-MTase activity of NSP14, we conducted thin-layer chromatography (TLC) analysis of NSP14 N7-MTase activity in the presence of C10^13^. Treatment with C10 inhibited NSP14-mediated methylation in a dose-dependent manner, as evidenced by the dramatic reduction in m7GpppG formation and accumulation of unmetabolized SAM and GpppG (**Extended Data Fig. 2d**). These results demonstrate that C10 is a potent inhibitor of NSP14 N7-MTase, providing a biochemical basis for its potential antiviral effects.

### C10 is a selective inhibitor of betacoronavirus NSP14

To evaluate whether C10 exhibits the broad-spectrum inhibitory activity against betacoronavirus NSP14, we assessed its inhibition of MERS-CoV NSP14 (**Extended Data Fig. 2e**), given the zoonotic risk it shares with SARS-CoV-2^46^. It revealed approximately two-fold lower potency compared to SARS-CoV-2 NSP14, exhibiting an IC_50_ value of 671.7 nM (**Fig. 1e**). We also prepared the NSP14 of HCoV-OC43, which belongs to the same β-coronavirus family as SARS-CoV-2^47^. C10 inhibited HCoV-OC43 NSP14 with approximately three-fold lower potency compared to SARS-CoV-2 NSP14, exhibiting an IC_50_ value of 1.02 µM in our biochemical assay (**Fig. 1f**). We next assessed whether C10 could effectively inhibit NSP14 from coronaviruses of different genera within the *Coronaviridae* family. We expressed and purified the NSP14 of the *Alphacoronavirus* HCoV-229E, which shares roughly 50% amino acid sequence identity with SARS-CoV-2 NSP14. The inhibitory activity of C10 against HCoV-229E NSP14 was approximately 30-fold reduced (IC_50_ = 9.1 µM) in our biochemical assays (**Fig. 1g**). Furthermore, it showed negligible inhibitory effects on Zika virus NS5 methyltransferase^48^ in the counter-screen (**Fig. 1h**). These findings confirm C10’s broad-spectrum inhibition against NSP14 of betacoronavirus.

To further test the target selectivity of C10, we tested the inhibitory effects of C10 against human methyltransferases, including m6A writer METTL3/14^49^, DNA m5C methyltransferase DNMT1^50^, and human mRNA cap guanine-N7 methyltransferase (hRNMT, **Extended Data Fig.2f**)^51^. In biochemical assays, C10 showed no detectable inhibitory activity against any of these enzymes (**Fig. 1i**). We also examined whether C-family compounds interfere with the ExoN activity of NSP14. Using an exonuclease assay with the purified NSP14-10 complex^52^, neither C3 nor C10 inhibited ExoN activity (**Fig. 1j, Extended Data Fig. 2g**). These results demonstrate that C-family compounds selectively inhibit the N7-MTase activity of NSP14 without affecting its ExoN function, thereby supporting their mechanism as highly specific methyltransferase inhibitors.

### Characterization of the inhibitory mode of lead compound

To elucidate the inhibitory mechanism of C10, we assessed its IC_50_ value as a function of both SAM and GpppG concentrations in the biochemical assay. Sinefungin acts as a positive control for a SAM-competitive inhibitor^53^. The IC_50_ value of sinefungin changed significantly from 1.59 µM to 52.90 µM with increasing SAM concentrations (**Fig. 2a**), while remaining constant around 1.59 µM across varying GpppG levels (**Fig. 2b**). Similar to sinefungin, the IC_50_ value of C10 showed a sharp shift from 0.34 µM to 5.70 µM in response to increased SAM concentration (**Fig. 2c**) but was not affected by GpppG concentration (**Fig. 2d**), demonstrating the SAM-competitive inhibition mechanism. This pattern was also observed in other C-family compounds, such as C3 and C7 (**Extended Data Fig. 3**), consistent with our initial virtual screening that intended to target the SAM-binding site of NSP14.

**Figure 2.**
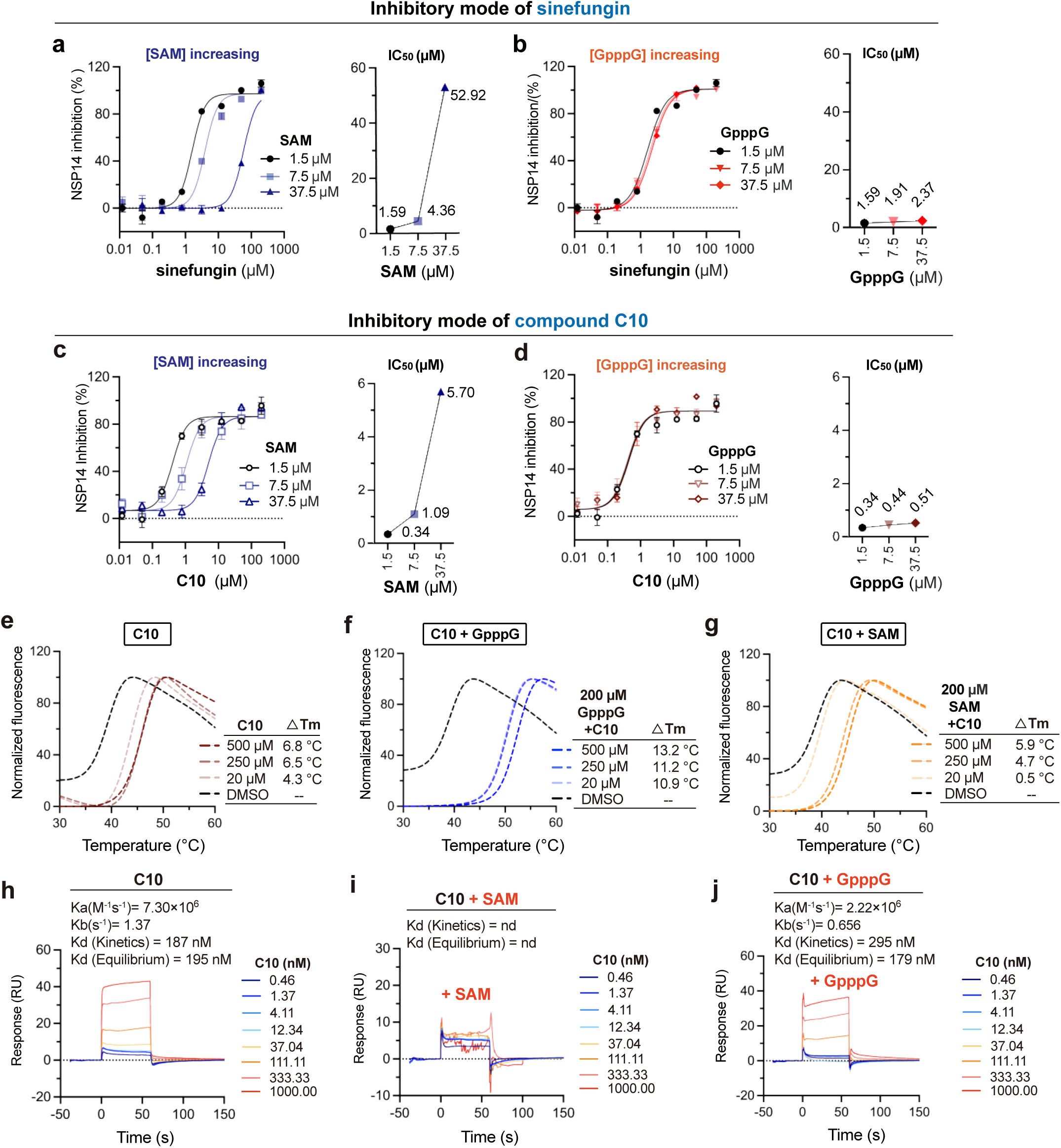
Characterization of the binding mode of C10 to SARS-CoV-2 NSP14. **a-b.** Dose-dependent inhibition curves of sinefungin against NSP14, with varying GpppG and SAM concentrations. SAM concentrations were increased while GpppG concentrations were fixed at 1.5 μM (**a**), or GpppG concentrations were increased while SAM concentrations were fixed at 1.5 μM (**b**). The obtained IC_50_ values are presented as scatter plots. Data are presented as mean ± SD from three independent replicates. **c-d.** Dose-dependent inhibition curves of C10 against NSP14 with varying concentrations of GpppG and SAM. In (c), SAM concentrations were increased while GpppG concentration was fixed at 1.5 μM. In (d), GpppG concentrations were increased while SAM concentrations were fixed at 1.5 μM. The IC_50_ values obtained from (c) and (d) are presented as scatter plots. Data are shown as mean ± SD from three independent replicates. **e.** Dose-dependent enhancement of the melting temperature (Tm) of SARS-CoV-2 NSP14 by compound C10 in the DSF assay. **f-g.** In the presence of 200 μM GpppG (f), instead of 200 μM SAM (g), treatment with C10 caused an additional increase in the Tm of NSP14, exhibiting a synergistic effect between C10 and GpppG. **h.** SPR binding data showing the interaction between C10 and immobilized NSP14, with data fitted to a 1:1 kinetic binding model. **i-j.** SPR binding data indicating that the binding affinity of C10 with immobilized NSP14 is reduced in the presence of SAM (i) compared to GpppG (j), suggesting a SAM-competitive binding mode.

We then used biophysical methods to confirm the binding mode of C10 to NSP14. In the differential scanning fluorimetry (DSF) assay, the addition of SAH induced a dose-dependent increase in the melting temperature (Tm) of NSP14, reaching up to a 10.0 °C shift, while SAM, GpppG, and m7GpppG caused only negligible Tm changes (**Extended Data Fig. 4a**). Treatment with C10 alone resulted in a dose-dependent increase in NSP14 stability, with ΔTm values of 4.3 °C at 20 μM, 6.5 °C at 250 μM, and 6.8 °C at 500 μM (**Fig. 2e, Extended Data Fig. 4b**), indicating direct binding of C10 to NSP14. Notably, co-incubation with 200 μM GpppG and increasing concentrations of C10 further enhanced Tm values, reaching 10.9 °C at 20 μM, 11.2 °C at 250 μM, and 13.2 °C at 500 μM (**Fig. 2f**). This synergistic stabilization effect was similarly observed with co-incubation of C10 and m7GpppG (**Extended Data Fig. 4c**), suggesting that C10 and GpppG/m7GpppG bind at distinct but adjacent sites within the catalytic pocket of NSP14. In contrast, co-incubation of C10 with 200 μM SAM or SAH did not show enhanced thermal stabilization (**Fig. 2g, Extended Data Fig. 4c)**, implying competition between C10 and SAM/SAH for the same or overlapping binding sites. The DSF data indicate that C10 specifically targets NSP14 in a SAM (and SAH)-competitive manner.

We next applied surface plasmon resonance (SPR) assay to determine the binding affinities and kinetic parameters of C-family compounds with NSP14 (**Extended Data Fig. 5**). The binding of C10 to NSP14 displayed an association rate (K_a_) of 7.3 × 10⁶ M⁻¹s⁻¹ and a dissociation rate (K_b_) of 1.37 s⁻¹, yielding a dissociation constant (K_d_) of 187 nM (**Fig. 2h**). When 20 μM SAM was added to the SPR running buffer, we observed that it led to undetectable binding between C10 and NSP14 (**Fig. 2i**). In contrast, the addition of 20 μM GpppG did not affect the binding of C10 to NSP14, as it still yielded a K_d_ value of 295 nM (**Fig. 2j**). A similar competition effect was observed for C10 in the presence of SAH, instead of m7GpppG (**Extended Data Fig. 6**). In summary, both biochemical and biophysical analyses demonstrate that C10 inhibits SARS-CoV-2 NSP14 by specifically occupying the SAM recognition pocket.

### The SAR understanding of compound C10

The crystal structures of NSP14 reported so far are predominantly in the SAH-bound form, emphasizing the essential role of SAH in stabilizing NSP14’s conformation, which is critical for efficient crystal packing. Our data demonstrate that compound C10 specifically occupies the SAH (and SAM)-binding site of NSP14. This interaction complicates obtaining a co-crystal structure of NSP14 bound to C10, but also provides valuable insights to guide our structural understanding of the C10-NSP14 binding mode via molecular docking and molecular dynamics (MD) simulations.

The simulations of SAH and m7GpppG in NSP14 catalytic pocket acted as the positive control, with stable RMSD fluctuation of SAH (**Extended Data Fig. 7a**) and binding details similar to reported crystal structures (PDB 7R2V for SAH, and 7QIF for m7GpppG) (**Extended Data Fig. 7b,c**). The RMSD fluctuation of bound C10 was observed (**Fig. 3a**), due to its dynamic purine ring (**Extended Data Fig. 7d**). The key residues for the binding of C10 to NSP14 predicted by the MM/GBSA free energy decomposition are listed (**Fig. 3b, Extended Data Fig. 7e).** The predicted structure of SARS-CoV-2 NSP14 bound to C10 and m7GpppG in a space-filling representation is shown in **Fig. 3c**. An expanded view of the bound C10 and m7GpppG (**Fig. 3d**) characterized a narrow binding pocket for C10, adjacent to the RNA cap-binding site. The core structure of C10 greatly overlaps with SAH (**Fig. 3e**), but the phenyl trifluoromethyl group (R1 part) of C10 extends out from the binding pocket of SAH (**Fig. 3f**). It was previously reported that an highly dynamic extended loop (residues 356–378) acts as the SAM entry gate of NSP14^27^. This SAM entry gate is in “open” state to embrace C10 in our predicted structure (**Fig. 3d**), while the closed conformation of this loop would not appear to be compatible with binding of C10 as it significantly overlaps with the core structure of C10 (**Extended Data Fig. 7f**). The enhanced potency of C10 relative to C1, driven by structural modifications to the R1 group, can be attributed to three main factors (**Fig. 3g**): Firstly, the ligand’s trifluoromethyl group inserts into a deep hydrophobic pocket lined by Gln313, Asp331 and Cys340. While amino group in sulfonamide forms hydrophobic interaction with Trp385. The stronger electronegativity of the phenyl trifluoromethyl group in C10 provides additional electrostatic interactions with residues of Site-C. Secondly, the amino group in sulfonamide of C10 forms a hydrogen bond with Gly333, and the amide group of the compound forms another hydrogen bond with Cys387. Finally, its aromatic ring in site C engages in p-π interaction with Arg310.

**Figure 3.**
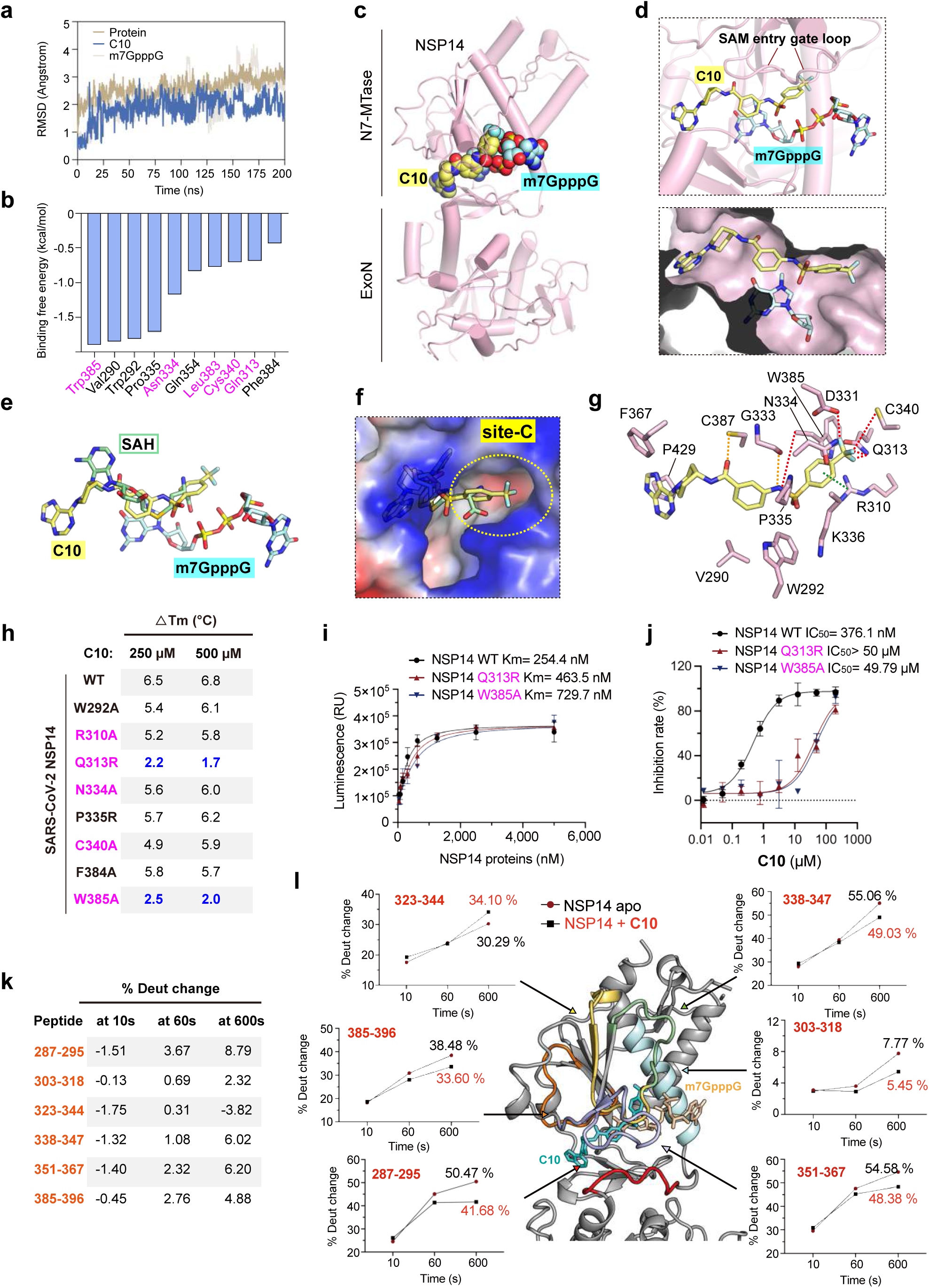
The structural understanding of the binding of compound C10 to NSP14. **a.** RMSD fluctuation of the C10-NSP14-m7GpppG complex within the simulation time. **b.** Top 10 key residues of NSP14 responsible for binding to C10 predicted by MM/GBSA. Residues of NSP14 site-C are colored magenta. **c.** Molecular docking structure of C10 and m7GpppG bound to NSP14. C10 and m7GpppG are shown in pale yellow and cyan, respectively, in sphere representations. NSP14 is shown in pink, cartoon representation. **d.** An expanded view of C10 and m7GpppG bound to the NSP14 catalytic pocket. C10 and m7GpppG are shown in pale yellow and cyan, respectively, in stick representations. NSP14 is shown in pink, cartoon (top) and surface (bottom) representation. **e.** Alignment of C10 (pale yellow) and SAH (cyan) bound to NSP14. **f.** C10 (pale yellow) extends from the SAH (cyan) -binding site to the adjacent site-C of NSP14, with protein shown in an electrostatic surface representation. Electrostatic surface potentials were calculated with PyMOL with thresholds ± 70. **g.** Key residues involved in the binding of C10 to NSP14. The red dashed lines represent hydrophobic interactions, the green dashed lines represent p-π conjugation, and the yellow dashed lines represent hydrogen bonding interactions. **h.** Summary of the ΔTm values for 250 μM and 500 μM C10 binding with SARS-CoV-2 NSP14 wild-type and mutants. **i.** Titration of NSP14 and mutants at 1.5 µM SAM and 1.5 µM GpppG for Km value determination. n = 3. Data are shown as mean ± SD. **j.** Dose-dependent inhibition curves of C10 against SARS-CoV-2 NSP14 wild-type, Q313R and W385A as measured by the MTase-Glo assay. **k.** The change of deuterium exchange rate is shown for six representative fragments after C10 binding at three time points corresponding to 10, 60 and 600 s. **l.** Regions of H/D exchange most affected by C10 binding to NSP14. The deuterium exchange rate is shown for six representative fragments in the presence (black squares) and absence (red circles) of C10. The black and red curves indicate the rate of H/D exchanges at three time points corresponding to 10, 60 and 600 s. Compound refers to C10.

To validate the key C10-binding sites on NSP14, we generated a series of single-point mutants (W292R, R310A, Q313R, N334A, P335R, C340A, F384A, and W385A) and assessed their interactions with C10 in DSF assays. Wild-type NSP14 exhibited a ΔTm of 6.5 °C and 6.8 °C with 250 and 500 µM C10 treatment (**Fig. 3h)**. In contrast, all mutants displayed reduced ΔTm values when incubating with C10, of which Q313R and W385A mutants, located in the defined site-C, showed the most significantly reduced ΔTm values of 2.2 and 2.5 °C, respectively (**Fig. 3h, Extended Data Fig. 8)**, indicating the critical role of these residues in C10 binding. We next evaluated the effects of Q313R and W385A mutations in the N7-MTase enzymatic assays. Interestingly, both Q313R and W385A mutants retained N7-MTase activity comparable to that of wild-type (WT) NSP14 (**Fig. 3i)**. However, both the Q313R mutation (IC_50_ > 50 µM) and the W385A mutation (IC_50_ = 49.79 µM) caused a drastic loss of C10’s inhibitory activity compared with wild-type NSP14 (IC_50_ = 376.1 nM) (**Fig. 3j)** These results indicate that Q313 and W385 are critical residues for C10 binding without affecting the intrinsic catalytic activity of NSP14.

To further gain structural insights into the C10-NSP14 interaction in the experimental end, we conducted hydrogen-deuterium exchange mass spectrometry (HDX-MS) analysis. Several peptide regions of NSP14 revealed significant changes in deuterium uptake upon C10 binding, as illustrated by differences in hydrogen-deuterium exchange rates at various time points (**Extended Data Fig. 9**). Peptides 385-396, 303-318, and 338-347, corresponding to Site-C, showed reduced deuterium incorporation by 4.88%, 2.32%, and 6.02%, respectively (**Fig. 3k,l**), suggesting interactions between the trifluoromethyl group of C10 and surrounding residues such as Gln313, Trp385, and Cys340. Peptide 287-295 exhibited the greatest reduction in deuterium uptake (8.79%), pointing to strong interactions with the sulfonamide moiety. Similarly, peptide 351-367, overlapping with the SAM/SAH interacting loop, showed a 6.02% reduction, consistent with binding of the pyrimidoimidazole ring. Interestingly, peptide 323–344 demonstrated increased deuterium exchange, likely reflecting localized conformational changes near the β-sheet region (**Fig. 3k,l**). Consistent with the molecular dynamics simulation results, the HDX-MS data provide experimental evidence for the predicted binding mode of C10 within the catalytic pocket of the NSP14 MTase domain.

### C10 exhibits potent anti-viral activity in cell-based assays

To verify whether C10 interacts with NSP14 in intact cells, we employed the cellular thermal shift assay (CETSA). HEK293T cells were transfected to overexpress SARS-CoV-2 NSP14 protein (HEK293-NSP14) and incubated with C10 for 8 hours at 40 hours post-transfection. Protein thermal stability was determined by western blot analysis after heating the cells^54^. Treatment with C10 at concentrations of 10 µM or 20 µM both led to a notable enhancement of the thermal stability of NSP14 (**Fig. 4a**). In addition, C10 treatment at concentrations ranging from 5 µM to 25 µM could dose-dependently stabilize NSP14 (**Fig. 4b**). These results clearly demonstrated the target engagement of C10 in live cells.

**Figure 4.**
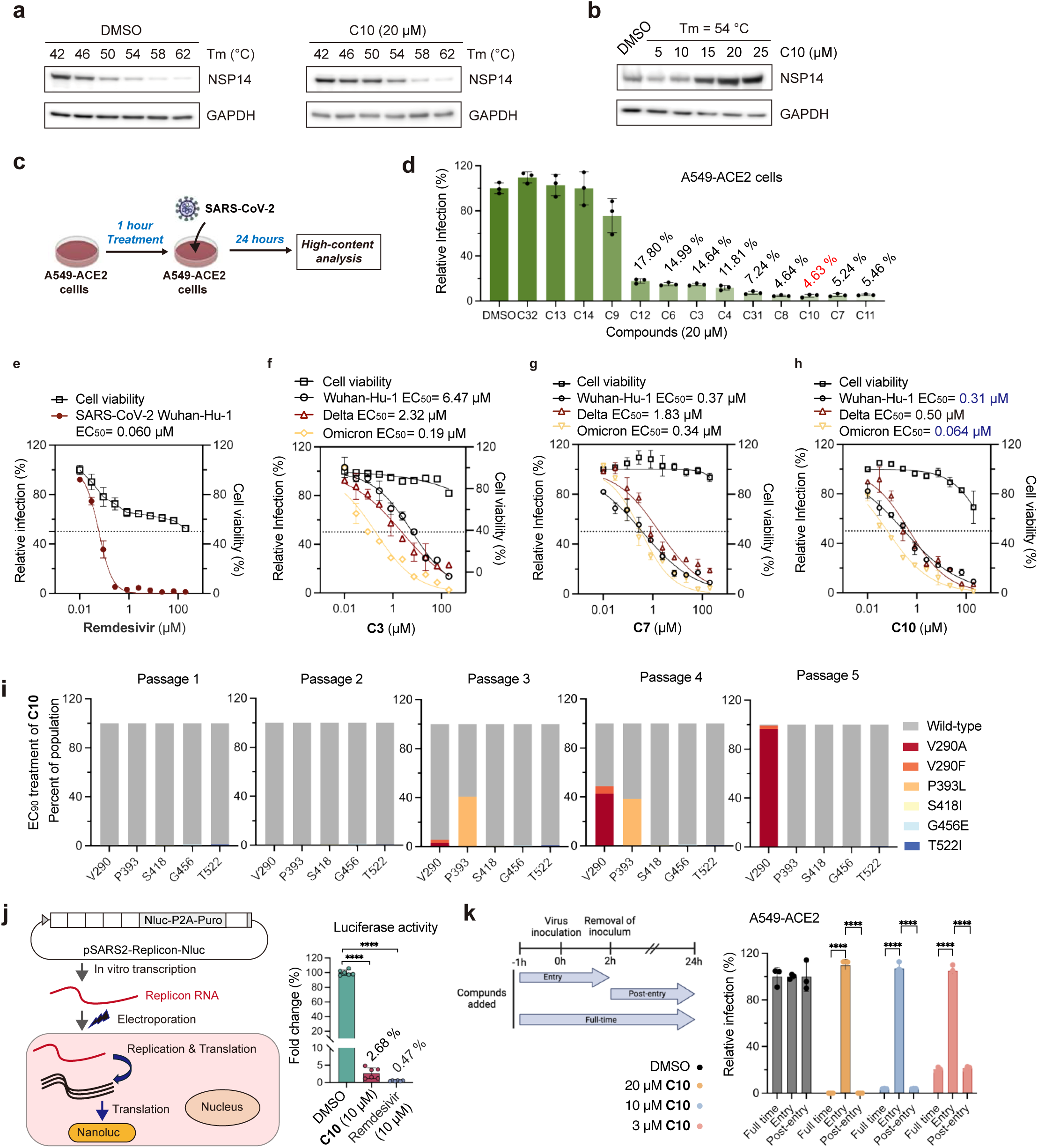
Evaluation of anti-viral activity in cell-based models. **a-b.** CETSA assays of HEK293-NSP14 cells treated with DMSO or 20 μM C10 at various heating temperatures (a), and with varying concentrations of C10 at 54 °C (b). **c.** Schematic diagram outlining the anti-viral activity assays conducted in cell-based assays. **d.** Histogram showing the infection ratios of A549-ACE2 cells treated with the selected C-family compounds at 20 μM. Data are presented as mean ± SD from three independent replicates. **e.** Dose-dependent inhibition curves of remdesivir against SARS-CoV-2 strain (Wuhan-Hu-1), along with the 50% cytotoxic concentration (CC_50_) values, as measured in A549-ACE2 cells. Data are presented as mean ± SD from three independent replicates. **f-h.** Dose-dependent inhibition curves of C3 (f), C7 (g), and C10 (h) against SARS-CoV-2 different strains in A549-ACE2 cells, with the 50% cytotoxic concentration (CC_50_) values. Data are presented as mean ± SD from three independent replicates. **i.** Frequency of NSP14 amino acid changes emerging in SARS-CoV-2 trVLP after 5 passages in Vero-N cells challenged with 10 μM or 20 μM C10. **j.** Schematic of the SARS-CoV-2 replicon assay (left) and quantification of replicon activity by luciferase reporter readout (right). Data are mean ± SD from three biological replicates. Statistical significance was determined by unpaired t-test; ****P < 0.0001. **k.** Time-of-addition analysis of C10. For all the experimental groups, cells were infected with SARS-CoV-2 trVLP at an MOI of 0.02, and the luciferase activity was measured. Data are presented as mean ± SD from three independent replicates. Statistical significance was determined by unpaired t-test, P****<0.0001.

To verify whether C-family compounds exhibit cellular activities, we performed cell-based antiviral assays. A549-ACE2 cells were infected with SARS-CoV-2 at an MOI of 0.2, and the percentage of infected cells was determined by immunofluorescence staining of SARS-CoV-2 nucleocapsid (N) protein and high-content imaging analysis at 24 hours post-infection (**Fig. 4c**). We first screened the potencies of C-family compounds at a concentration of 20 µM. We observed a reduced percentage of infected cells treated by C-family compounds, with up to a 20-fold reduction in SARS-CoV-2 infection, compared to the untreated cells (**Fig. 4d**). We next selected C10 and other top-performing compounds to determine their dose-response inhibitory curves against SARS-CoV-2 original strain and other variants (**Extended Data Fig. 10a**). FDA-approved drug remdesivir, a well-known anti-SARS-CoV-2 drug by inhibiting viral RNA synthesis^55,56^, served as the positive control by exhibiting the half-maximal effective concentration (EC_50_) value of 60 nM in our cell-based assays (**Fig. 4e**). Top-performing compounds C3, C7, and C10 potently reduced the infectivity of original SARS-CoV-2 and its variants (**Fig. 4f-h**). Notably, the potency of C10 in inhibiting original SARS-CoV-2 and Delta variant with EC_50_ values of 0.31 µM and 0.50 µM, respectively, and inhibiting the Omicron variant reached up to 64 nM with low cytotoxicity (CC_50_ > 100 µM, **Fig. 4h**), displaying a high selectivity index (SI > 1500). C10 inhibited HCoV-OC43 infection in A549-ACE2 cells with an EC_50_ value of 4.36 µM, while the inhibition of HCoV-229E infection was negligible (**Extended Data Fig. 10b,c**). We conclude that C10 holds promise as a lead candidate for developing effective treatments against SARS-CoV-2 and emerging variants.

### Resistance mutations to compound C10

To identify resistance mutations and further elucidate the target site of compound C10, we conducted serial passaging under increasing concentrations of C10 by using the SARS-CoV-2 trVLP system, which consists of N gene-deleted genomic viral RNA and Vero-N cells^57–59^. This system enables completion of the full viral life cycle exclusively in cells expressing the SARS-CoV-2 N protein. After five passages, we observed a gradual reduction in C10’s antiviral efficacy (**Extended Data Fig. 11a,b**), indicating the emergence of drug-resistant variants. Next-generation sequencing (NGS) of the passaged viral populations revealed no significant mutations in NSP14 during the first two passages. However, by the third and fourth passages, we detected the emergence of V290A and P393L mutations in NSP14 at high frequencies. Notably, by the fifth passage, the V290A mutation reached a frequency of 96.73%, suggesting strong positive selection under C10 pressure (**Fig. 4i, Extended Data Fig. 11c)**.

We next performed fitness competition assays by mixing wild-type and V290A SARS-CoV-2 trVLP at ratios of 10:0, 0:10, and 9:1, and passaging them for two rounds. NGS analysis showed that in the 9:1 mixture, the wild-type NSP14 rapidly outcompeted the V290A mutant, with its proportion increasing to 94.34% after first passage, and reaching 100% after the two passages (**Extended Data Fig. 11d**). This suggests that the V290A mutation, although conferring resistance under drug pressure, reduces viral fitness in the absence of C10.

To validate the functional relevance of identified resistant mutations, NSP14 proteins bearing the V290A or P393L mutations were measured bindings to C10 in DSF assay. V290A and P393L mutants exhibited significantly reduced thermal stability (ΔTm=0.9 and 5.8°C) upon 250 µM C10 binding (**Extended Data Fig. 11e,f**), compared to wild-type NSP14 (ΔTm=6.5 °C), indicating the critical role of these residues in C10 binding. Notably, the V290A mutation is consistent with our molecular simulation results, which highlight its direct involvement in the C10 binding interface (**Fig.3b**). In contrast, the P393L mutation resides outside the catalytic pocket and is embedded within the structural core of NSP14. We therefore hypothesize that P393L may influence C10 binding through an allosteric mechanism.

### C10 inhibits SARS-CoV-2 at the replication stage

To examine whether C10 inhibits SARS-CoV-2 at the replication stage, we utilized the SARS-CoV-2 replicon system by replacing the spike to ORF8 region with a Nanoluciferase-P2A-puromycin cassette reporter^60^ (**Fig. 4j, left**). HeLa cells were electroporated with *in vitro*-transcribed and capped replicon RNA and then incubated with 10 µM remdesivir (the positive control) or 10 µM C10. Replication inhibition was determined by the reduction of luciferase activity relative to that in DMSO control-treated cells at 24 hours post-electroporation. Both compounds dramatically reduced the replication levels of SARS-CoV-2, as the relative luciferase signals in cells treated with remdesivir and C10 were about 2.68% and 0.47%, respectively, lower than that in DMSO-treated cells (**Fig. 4j, right**). These data indicate that C10 is a potent antiviral compound against the replication of SARS-CoV-2.

We next performed time-of-addition (TOA) experiments to further elucidate the mechanism of action of C10, using SARS-CoV-2 trVLP system in A549-ACE2 cells. C10 was treated at concentrations of 3 μM, 10 μM, and 20 μM during different stages of the viral life cycle: full-time treatment, entry phase, and post-entry phase (**Fig. 4k, left**). Our results show that C10 exhibits significantly stronger inhibitory activity during full-time and post-entry treatments, while minimal inhibition was observed when applied only during the entry phase (**Fig. 4k, right**). These findings suggest that C10 primarily targets the viral replication stage, consistent with its proposed mechanism of action as an NSP14 inhibitor.

### C10 suppresses viral translation and exhibits immunostimulatory effect

N7 methylation of the mRNA cap is critical for optimal recognition by eukaryotic translation initiation factors, thereby supporting the translation of host and viral mRNAs. We performed polysome profiles of mock-infected and VLP-infected cell lysates, which indicates that SARS-CoV-2 trVLP infection slightly downregulated host translation (**Extended Data Fig. 12**). To further evaluate the impact of C10 on mRNA translation, we pre-treated A549-ACE2 cells with DMSO, 20 μM, or 40 μM C10, followed by infection with SARS-CoV-2 trVLP for 8 hours. Polysome profiling of cell lysates showed comparable overall profiles in the presence or absence of C10 (**Fig. 5a**), indicating no global disruption of host translation. To investigate whether C10 selectively impairs viral translation, we assessed the abundance of SARS-CoV-2 Spike mRNA relative to host HPRT1 mRNA in different polysome fractions by RT-qPCR. Treatment with 20 μM and 40 μM C10 resulted in a 10-fold and 13-fold reduction respectively, compared to DMSO prior to polysome fractionation (**Fig. 5b, left**). Notably, this reduction was more significant in the heavy polysome fractions (∼13-fold and ∼40-fold for 20 μM and 40 μM C10, respectively) compared to the free polysome fractions (∼2.5-fold and ∼26-fold) (**Fig. 5b, right**), indicating impaired ribosome engagement with viral mRNA. These findings suggest that C10, by inhibiting NSP14 MTase activity, disrupts mRNA capping, thereby selectively suppressing viral translation without broadly affecting host protein synthesis.

**Figure 5.**
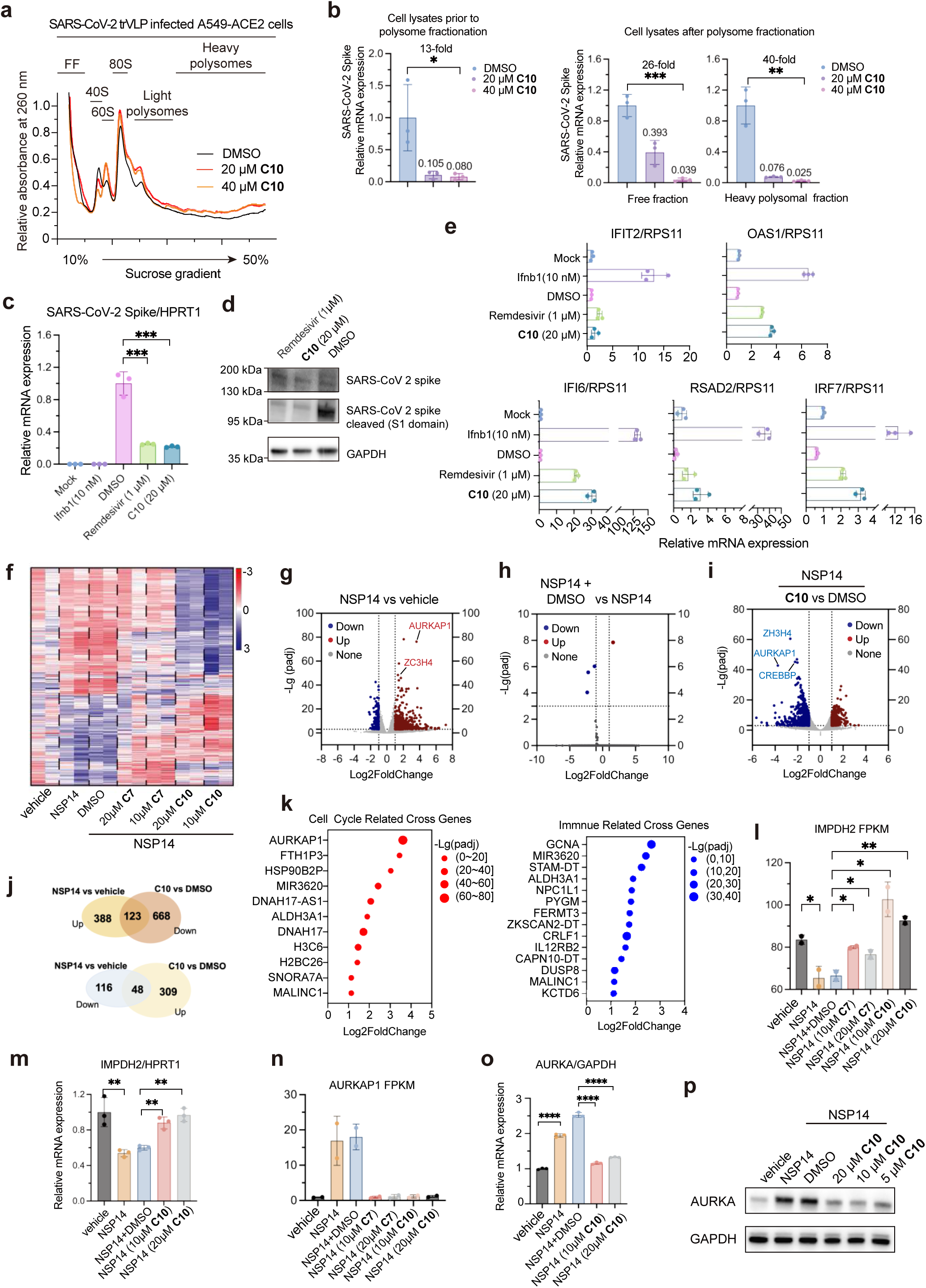
C10 suppresses viral translation and modulates host responses. **a.** Polysome profiling of SARS-CoV-2 trVLP infected A549-ACE2 cells treated with C10 (20 µM or 40 µM) or DMSO for 8 h, respectively. FF, free fraction. **b.** Left panel: Ratio of SARS-CoV-2 Spike mRNA relative to host *HPRT1* in lysates of A549-ACE2 cells treated with C10 (20 µM or 40 µM) or DMSO and infected with SARS-CoV-2 trVLP for 8 h, respectively, prior to polysome fractionation. Right panel: Ratio of SARS-CoV-2 trVLP Spike mRNA relative to host *HPRT1* in the free fraction or heavy polysomal fraction of lysates of A549-ACE2 cells treated with C10 (20 µM or 40 µM) or DMSO and infected with SARS-CoV-2 trVLP for 8 h, respectively, after polysome fractionation. n = 3; mean ± SD. Adjusted P values by ordinary one-way ANOVA are given. P * < 0.05; P ** <0.01; P *** <0.001. **c-d.** Quantification of Spike expression by RT-qPCR (c) and western blotting (d) in SARS-CoV-2 trVLP infected A549-ACE2 cells treated with 20 µM C10. **e.** C10 inhibition of SARS-CoV-2 trVLP replication induces ISG expression in A549-ACE2 cells. Cells were mock-infected or infected with SARS-CoV-2 trVLP, pre-treated for 1 h with DMSO, C10 (20 µM), or remdesivir (1 µM), respectively; IFNB1 protein (10 nM) served as a positive control. Host and viral transcripts were quantified by RT-qPCR 24 h post-infection. Data are mean values from *n* = 3 biological replicates. **f.** Cluster heatmap showing differentially expressed transcription factors from RNA-seq data. Data are presented as averages of two independent experiments. **g-i.** Volcano plots of RNA-seq data displaying upregulated and downregulated gene expression (2-fold change, FDR ≤ 0.001) in HEK293T-NSP14 cells compared to vehicle (g), DMSO treated compared to untreated HEK293T-NSP14 cells (h), and C10 (20 μM) treated compared to DMSO treated HEK293-NSP14 cells (i). *P*.adj refers to the adjusted P value. Data are presented as averages of two independent experiments. **j.** Venn diagram illustrating the overlap of upregulated and downregulated genes between NSP14 vs vehicle and 20 μM C10 vs DMSO conditions. Data are presented as averages of two independent experiments. **k.** Bubble plots showing cross-regulated genes that are upregulated by NSP14 expression and downregulated by C10 in HEK293T-NSP14 cells. Left: Cell cycle-related cross-regulated genes. Right: Immune-related cross-regulated genes. **l.** FPKM analysis of *IMPDH2* from RNA-seq data. Data are presented as averages of two independent experiments. **m.** *IMPDH2* transcriptional changes as detected by qPCR. Data are presented as mean ± SD from three replicates. Statistical significance was analyzed by unpaired t-test. P****<0.0001. **n.** FPKM analysis of *AURKAP1* from RNA-seq data. Data are presented as averages of two independent experiments. **o.** *AURKA* transcriptional changes as detected by qPCR. Data are presented as mean ± SD from three replicates. Statistical significance was analyzed by unpaired t-test. P****<0.0001. **p.** Protein expression changes of AURKA detected by Western blot.

To explore the immunostimulatory effects of C10 treatment, we measured the expression of several interferon-stimulated genes (ISGs)^44,61^, including RSAD2, IFI6, IRF7, IFIT2, and OAS1, in SARS-CoV-2 trVLP and A549-ACE2 cells system. In parallel, qPCR and western blot analyses confirmed that 20 μM C10 significantly reduced mRNA and protein expression of SARS-CoV-2 Spike, with remdesivir as a positive control (**Fig. 5c,d**). When compared to DMSO or Remdesivir, C10 treatment at 20 μM led to a modest increase in ISG expression (**Fig. 5e**), which is consistent with previously reported results of NSP14 inhibition by TDI-015051^44^. These findings suggest that C10’s inhibition of RNA capping triggers an immunostimulatory effect, which appears secondary to its primary antiviral effect. This immunological consequence likely contributes to the enhanced viral clearance observed with C10 treatment.

### C10 reverses NSP14-mediated host transcriptome modulation

Among all SARS-CoV-2 non-structural proteins, NSP14 has been uniquely shown to induce extensive remodeling of the host transcriptome, closely resembling the alterations observed during SARS-CoV-2 infection^15^. Additionally, NSP14 expression led to changes in the splicing of over 1,000 genes, implicating a pathogenic role that extends beyond its canonical function in viral RNA capping. Notably, these transcriptomic and splicing effects are independent of the ExoN domain but require an intact N7-MTase domain^15^. We infer that small-molecule inhibition of NSP14 methyltransferase, as achieved by C10, could reverse NSP14-driven host transcriptome modulation.

To test this, we performed RNA sequencing (RNA-seq) in HEK293T cells transfected with an NSP14 expression plasmid containing EGFP reporter, followed by treatment with either DMSO or C10^15^. Heatmap clustering of differentially expressed transcription factors revealed that C10 specifically reversed NSP14-induced transcriptional changes (**Fig. 5f**). Consistent with previous reports, NSP14 expression significantly altered the expression of 675 genes (511 upregulated, 164 downregulated; ≥2-fold change, FDR ≤ 0.001) (**Fig. 5g**). In contrast, DMSO treatment minimally affected gene expression in NSP14-expressing cells (5 DEGs total; **Fig. 5h**). Notably, treatment with 20 μM C10 in NSP14-expressing cells altered 1,168 genes (357 downregulated, 811 downregulated; ≥2-fold change, FDR ≤ 0.001) (**Fig. 5i**). Venn diagram analysis revealed that the major overlap between NSP14- and C10-induced DEGs reflected opposing regulatory effects. Genes upregulated by NSP14 were frequently downregulated by C10, and vice versa (**Fig. 5j**). Pathway enrichment analysis showed that many C10-regulated genes are involved in cell cycle control and immune-related pathways (**Fig. 5k, Extended Data Fig. 13a**). Another lead compound, C7, produced a similar reversal effect, confirming that C-family inhibitors specifically target NSP14-mediated host transcriptome modulation (**Extended Data Fig. 14**).

Several of the most enriched genes overlapped with those reported in the previous study, such as *CREBBP* and *Z3CH4*^15^ **(Extended Data Fig. 13b, c**). Guanosine triphosphate (GTP) levels inversely regulate inosine monophosphate dehydrogenase 2 (IMPDH2) expression, and NSP14 has been shown to activate IMPDH2, elevating GTP levels and thereby suppressing *IMPDH2* mRNA expression^23,62^. Consistent with this mechanism, our RNA-seq and RT-qPCR analyses confirmed that NSP14 expression reduced *IMPDH2* transcript levels (**Fig. 5l,m**). C10 treatment dose-dependently restored *IMPDH2* mRNA levels (**Fig. 5l,m**), supporting the conclusion that C10 counteracts NSP14-mediated downregulation of IMPDH2.

Beyond these known associations, our data identified novel host factors linked to NSP14 activity. In our RNA-seq dataset, *AURKAP1* (Aurora kinase A pseudogene 1) was among the most enriched transcripts in NSP14-expressing cells, and its enrichment was reversed by C10 (**Fig. 5n**). Proteomic profiling of SARS-CoV-2 infected cells has highlighted cell cycle regulators such as Aurora kinase A (AURKA) as highly enriched host proteins^63^. Given that pseudogenes can regulate their parental genes, we examined *AURKA* itself and found that NSP14 markedly increased both *AURKA* mRNA and protein levels, while C10 treatment dose-dependently reduced *AURKA* expression at both transcript and protein levels (**Fig. 5o,p**)., and thus potentially driving host cell cycle arrest.

To test whether NSP14 could induce cell cycle arrest at the phenotypic level, we performed flow cytometry to analyze the effects of NSP14 on the cell cycle distribution. It revealed that 4.04% of cells in the vehicle group were in the G2/M phase, while 25.52% of cells were in the G2/M phase after NSP14 expression (**Extended Data Fig. 15a**), showing a clear NSP14-induced accumulation of cells in the G2/M phase, which was consistently observed in SARS-CoV-2-infected cells^63^. We next test whether C10-mediated inhibition of NSP14 could potentially reverse NSP14-induced cell cycle arrest (**Extended Data Fig. 15a**). As expected, 8.61% of cells were in the G2/M phase in the 10 µM C10-treated group, and 4.88% of cells were in the G2/M phase in the 20 µM C10-treated group (**Extended Data Fig. 15b**), suggesting a dose-dependent recovery from cell cycle arrest. Our data uncover the unexpected benefit of NSP14 inhibition in protecting host cells from cell cycle arrest during SARS-CoV-2 infection.

### In vivo antiviral efficacy of C10

The lead compounds C7 and C10 were selected for 24-hour pharmacokinetic (PK) studies in mice to assess their in vivo bioavailability. Three male ICR mice per group were dosed with the compounds at 50 mg/kg and 70 mg/kg via intraperitoneal injection (I.P.). Blood samples were collected at 0.25, 0.5, 1, 2, 4, 8, 12, and 24 hours, and the drug concentrations were quantified by LC-MS/MS. C10 exhibited faster absorption and higher plasma concentrations in vivo compared to C7 (**Fig. 6a** and **Extended Data Fig. 16**). C10 was rapidly absorbed after I.P. at 70 mg/kg and reached the maximum plasma concentration (C_max_) at 1.0 hour (T_max_), with a peak concentration of 19,450.11 ng/mL. This resulted in favorable plasma exposure, with an area under the curve (AUC) of 37,779.33 ng/mL (**Fig. 6b**). C10 exhibited moderate clearance, with a half-life (T_1/2_) of 1.71 hours. Plasma drug concentrations remained above the antiviral EC_50_ value (0.31 μM) for 6 hours. Next, young C57BL/6 uninfected mice were intraperitoneally injected with C10 at doses of 300 mg/kg, 200 mg/kg, and 100 mg/kg twice daily (B.I.D.) for 3 days. The weight loss plot shows that mice treated with 300 mg/kg C10 experienced rapid body weight loss exceeding 20% (**Fig. 6c**), leading to a 100% fatality rate by 3 days post-inoculation (D.P.I.) due to drug toxicity (**Fig. 6d**). In contrast, most mice treated with 100 mg/kg and 200 mg/kg C10 exhibited reduced weight loss of less than 10% and 20%, respectively, leading to improved survival rates (100%, P < 0.0001) (**Fig. 6c** and **6d**).

**Figure 6.**
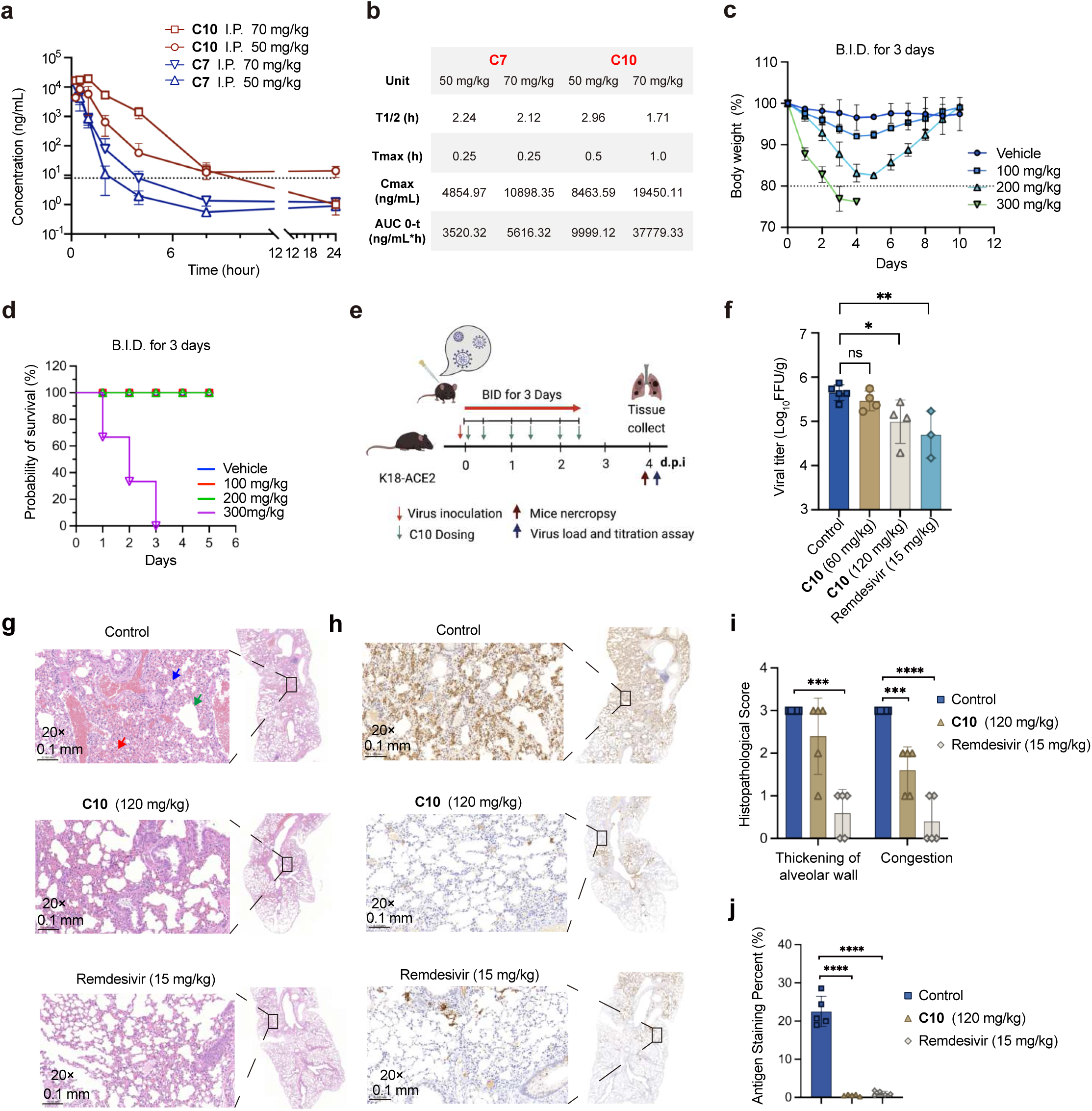
Antiviral efficacy of C10 in a K18-hACE2 transgenic mouse model. **a.** Plasma drug concentrations of C7 and C10 in male ICR mice (6-8 weeks) following intraperitoneal injection of 50 mg/kg (C7) and 70 mg/kg (C10) in a 5% dimethyl sulfoxide and 95% cyclodextrin solution (m(cyclodextrin):m(normal saline) = 1:2) (n = 3 per group). I.P. denotes intraperitoneal injection. **b.** In vitro pharmacokinetic (PK) parameters of C7 and C10. **c-d**. Body weight loss (c) and survival rate (d) without virus infection. BID: twice a day. **e.** Experimental design for the study of 3-day treatment study. K18-ACE2 mice were intranasally (IN) inoculated with 5000 PFU of SARS-CoV-2 original strain, followed by intraperitoneal injection of 60 mg/kg, 120 mg/kg, or 180 mg/kg C10, 15 mg/kg remdesivir, or vehicle twice a day (BID) for 3 days. **f.** Viral lung titers in the lungs of mice in all study groups were determined on day 4 post-infection. Lung titers are presented as log10 PFU/g of lung tissue. Adjusted P values from ordinary one-way analysis of variance are indicated. Sample sizes: n = 5 for group 1, n = 4 for groups 2 and 3, and n = 3 for group 4. Data are shown as mean ± SD, analyzed using unpaired t-test with Welch’s correction. *P < 0.05. **g.** Lungs collected at 4 DPI from vehicle, 120 mg/kg C10 and 15 mg/kg Remdesivir treated mice were stained with haematoxylin and eosin (H&E). H&E stained lungs from vehicle-treated infected mice exhibited severe thickening of alveolar wall (blue arrow), congestion of alveolar wall capillaries (red arrow), and slight dilation of alveoli (green arrow). Scale bars, 0.100 mm. **h.** Lungs from vehicle-treated, C10-treated and Remdesivir-treated mice were immunostained to detect SARS-CoV-2 nucleocapsid protein (brown color staining). Scale bars, 0.100 mm. **i-j**. Summary scores of lung lesions (i) and of nucleocapsid immunostaining of lungs (j). Data are shown as mean ± SD and analyzed with unpaired t-test with Welch’s correction. Ns, not significant; P ** <0.01; P *** <0.001; P**** < 0.0001.

To assess the antiviral efficacy of C10 in vivo, we used the K18-hACE2 transgenic mouse model. The K18-ACE2 mice (6 to 8 weeks old) were intranasally inoculated with SARS-CoV-2 original strain^64,65^. We then evaluated the efficacy of I.P. C10 in reducing SARS-CoV-2 viral burden in the lungs of this mouse model, with remdesivir as the control. Inhibitors and vehicle controls were dosed by I.P. at one hour post-infection, followed by B.I.D. dosing from day 0 to day 3 post-infection (**Fig. 6e**). The viral load in the lungs was determined at day 4 post-infection and revealed that vehicle-treated mice have robust infections in the lungs, with a mean lung titer of Log10 PFU/g = 5.73. In contrast, 120 mg/kg C10-treated mice had significantly lower lung viral titers (mean titer of Log10 PFU/g = 4.99, P < 0.05) (**Fig. 6f**). Mice treated with 60 mg/kg C10 showed reduced lung viral titers compared to vehicle-treated controls, though the difference was not statistically significant (**Fig. 6f**). These results highlight the need for further structural optimization of these inhibitors.

Histopathological analysis of the lungs from vehicle-treated mice infected with the SARS-CoV-2 nCoV original strain at 4 dpi revealed multifocal pulmonary lesions, including congestion, alveolar wall thickening, and inflammatory infiltration (**Fig. 6g**). In contrast, these pathological features were markedly alleviated in mice treated with 120 mg/kg C10 or 15 mg/kg Remdesivir (**Fig. 6g**). Immunohistochemical analysis using a monoclonal antibody against the SARS-CoV-2 N protein demonstrated strong and widespread antigen staining in the lungs of vehicle-treated mice (**Fig. 6h**). However, treatment with C10 or Remdesivir significantly reduced viral antigen staining, with only a few sporadic positive cells observed (**Fig. 6h**). The pathological lesions and immunostaining were quantified (**Fig. 6i** and **6j**), respectively, which are in line with the lung viral titer results (**Fig. 6f**).

## Discussion

Our study demonstrates that C10 is a potent and selective NSP14 N7-methyltransferase inhibitor with potent antiviral effects against SARS-CoV-2 and its variants. Structure-based virtual screening followed by rational design and lead optimization enabled the identification of compound C10 featuring a unique structural scaffold, which binds specifically to the SAM-binding pocket of NSP14. The structural modifications leading to C10 enhanced both its potency and selectivity, as evidenced by its nanomolar EC_50_ values across multiple SARS-CoV-2 variants and its ability to inhibit β-coronavirus NSP14 with significantly higher specificity compared to other viral and human methyltransferases.

The favorable toxicity profile of C10, demonstrated in both cell-based and murine models, is attributable to its superior target selectivity. This pharmacological advantage arises from C10’s precise molecular recognition of viral-specific structural motifs within the N7-MTase domain, while effectively avoiding interference with host methyltransferase. Guided by orthogonal biochemical and biophysical data, as well as molecular dynamics simulations and HDX-MS, we identified one of the key determinants of C10’s selectivity: its R1 group binds to a unique druggable site-C within the catalytic pocket of NSP14, positioned adjacent to the SAM and RNA cap binding sites. The trifluoromethyl-phenyl moiety of C10 engages in critical stacking interactions with Arg310, and forms stabilizing electrostatic interactions in the site-C pocket. These structural insights reveal how C10 achieves high affinity and selectivity, distinguishing it from nonspecific SAM competitors. The engagement of site-C is a novel observation, offering an opportunity for structure-guided drug design. Our results identify V290 and P393 as critical residues involved in C10 binding to NSP14 and reveal a resistance mechanism driven by the V290A substitution. Although the V290A mutation confers resistance, its reduced viral fitness highlights a potential vulnerability that could be leveraged in combination therapies or for the development of improved analogs.

Our current findings provide preliminary proof-of-concept that NSP14 inhibition exhibits multiple antiviral effects. However, the effective dose of C10 is approximately eight-fold higher than that of remdesivir, indicating substantial medicinal chemical modification will be required to improve the therapeutic index before clinical translation. Future medicinal chemistry efforts will focus on further optimizing the pharmacokinetic properties of C10 while maintaining its high potency and selectivity. Additionally, TDI-015051, another first-in-class inhibitor of the NSP14 RNA cap binding pocket, occupies the GpppG binding site and forming cooperative interactions with SAH. This inspires structural optimization of C10, which could potentially target the RNA cap substrate or act as bi-substrate inhibitors that engage both the SAM and RNA substrate binding sites^34^. These SAR insights, as gained from C10 and TDI-015051, will also guide the design of next-generation NSP14 inhibitors, potentially broadening their applicability to other viral methyltransferases. Given that mRNA cap methyltransferases are conserved across multiple RNA and DNA viruses, targeting these enzymes with structurally optimized inhibitors like C10 may offer a new avenue for developing broad-spectrum antiviral therapeutics.

Beyond its direct antiviral activity, C10 also modulates host cellular pathways perturbed by SARS-CoV-2 infection. Our transcriptomic and phenotypic analyses revealed that NSP14 induces host cell cycle arrest by upregulating key regulators such as AURKA, a previously unrecognized function of this viral protein. C10 treatment effectively C10 specifically reversed NSP14-mediated alterations in host transcriptome, rescuing normal cell cycle progression disrupted by NSP14. This dual mechanism, direct viral inhibition and modulation of host-virus interactions, further strengthens the therapeutic potential of C10. These pharmacological mechanisms could also be shared by RNA capping methyltransferases from other pandemic-related viruses, such as Zika virus and Mpox virus, thereby position RNA capping as a prime target for new antiviral drug discovery opportunities.

## Author contribution

W.X., X.D., and R.Z. conceptualized and supervised the project. M.L. conducted the screening, biochemical and biophysical assays, RNA sequencing, and cell cycle analysis, in which H.W., K.C., Y.C., W.C., X.L., H.Z. and B.L. provided experimental support. M.J. and J.C. designed and synthesized the inhibitors. Z.W. and P.Z. performed antiviral evaluations in both cell-based and mouse models. L.S. and L.L. carried out virtual screening, while J.W. and K.X. performed molecular dynamics simulations. X.W., under the supervision of Y.X., helped the polysome profiling experiments. F.X. and Q.Z. contributed intellectual expertise and key methodologies. W.X. and M.L. drafted the manuscript, with contributions from all authors. All authors reviewed and approved the final version.

## Declaration of generative AI and AI-assisted technologies in the writing process

During the preparation of this work ChatGPT 4.0 was used in order to check spelling and to correct the grammar. After using this tool, the authors reviewed and edited the content and take full responsibility for the content of the publication.

## Data availability

All data are available from the corresponding authors upon reasonable request. RNA seq data are available under GEO accession numbers GSE290079. NGS data are available under GEO accession numbers GSE304814.

## Declaration of competing interest

X.D., W.X., J.M., M.L., K.C., and J.C. are listed as inventors on a provisional patent that relates to the methods and compounds described here. The other authors declare no competing interests.

## Acknowledgment

We thank Jie Ma from the core facilities of the Life Sciences Institute Zhejiang University for SPR support. We thank the core facilities of Zhejiang University School of Medicine for technical support. We thank Feng Xu and Weiyuan Jin at the Westlake University High-Throughput Core Facility. We thank the Institutional Technology Service Center of Shanghai Institute of Materia Medica, Chinese Academy of Sciences (Shanghai, China) for technical assistance in HDX-MS experiments and analysis. This research was funded by the National Natural Science Foundation of China (Grant 82151216 and 82473975 to W. Xie, Grant 82341084 and 32270163 to R. Zhang), Shanghai Municipal Science and Technology Major Project (ZD2021CY001), Non-profit Central Research Institute Fund of Chinese Academy of Medical Sciences (2023-PT310-02).

## Materials and Methods

### Virtual screening

The X-ray co-crystallized structure of the SARS-CoV-2 NSP14 protein (PDB ID: 7R2V) was retrieved from the RCSB-PDB database and processed using the Protein Preparation Workflow (Schrödinger Suite 2021-2) in Maestro (version 12.8). Briefly, the default protocols were employed to refine the structure by assigning bond orders, adding hydrogens, creating zero-order bonds for metals, disulfide bonds, and filling in missing side chains using Prime. Heterogeneous states were generated with the Epik program (version 5.6) at pH 7.4 ± 2.0. Hydrogen bond assignments were optimized by sampling water orientations and using PROPKA at pH 7.4. Finally, restrained energy minimization was performed using the OPLS4 force field, ensuring the convergence of heavy atoms to an RMSD of 0.30 Å. All ligands used for molecular docking were prepared using the default parameters in LigPrep (Schrödinger Suite 2021-2) within Maestro (version 12.8). The preparation process included hydrogenation, salt removal, tautomer generation, and the calculation of ionization states, performed with the Epik program (version 5.6) at pH 7.0 ± 2.0 under the OPLS4 force field. Additionally, up to 32 stereoisomers were generated for each ligand, maintaining specific chirality under the defined computational conditions. The protein’s active pocket, with a diameter of 20 Å, based on the binding site of the original ligand, was constructed using the Receptor Grid Generation (Schrödinger Suite 2021-2). Default parameters, such as Vander Waals radius scaling factor 1.0 and partial charge cutoff 0.25, are utilized in this procedure. lt is worth noting that the OPLS4 forcefield was applied instead of normal OPLS 2005.

Molecular docking was performed using the Glide (version 9.1) program. The ligands were sampled in a flexible pattern, including sampling nitrogen inversion, sampling ring conformations as well as bias sampling of torsions for all predefined functional groups. Followed by the standard precision (SP) algorithm and the docking pocket identified by the above method, all the prepared ligands were docked under the OPLS4 forcefield, except for molecules with an atomic number of more than 500 or a heavy atomic number of more than 100. The SP mode employs basic molecular mechanics force fields and simplified scoring functions, incorporating limited ligand conformational sampling while maintaining receptor rigidity, making it suitable for rapid filtration in large-scale virtual screening. A maximum of ten poses per ligand were written out, all of which were subjected to post-docking minimization. Docking score, which is a Glide score with Epik state penalties, was employed as a scoring function to rank-order compounds. Afterward, ligands with a docking score better than -6 kcal/mol were incorporated into the next screening, which adopted the extra precision (XP) algorithm. The XP mode integrates hybrid QM/MM methods with refined scoring functions, enabling extensive conformational sampling including receptor key residue flexibility through rotamer library sampling, along with explicit solvation models to provide more accurate binding mode predictions during lead optimization stages. The parameter settings of the XP algorithm were the same as those of the SP algorithm. Eventually, molecules with a docking score better than -9 kcal/mol from XP were selected for the next step. The DeepDock program, available on GitHub, is a deep learning framework for protein-ligand docking, designed to predict binding affinities between proteins and small molecules. It utilizes a convolutional neural network (CNN) to learn the relationship between the three-dimensional structures of proteins and ligands and their corresponding binding affinities. In our research, we applied the MaSIF algorithm to analyze the interfacial characteristics of NSP14 when bound to the molecule. The software was subsequently used to perform docking and scoring of various molecules.

### Protein expression and purification

Codon-optimized SARS-CoV-2 NSP14 was synthesized by Tsingke Biotechnology and inserted into the pET-28a vector (Novagen) with an N-terminal His6-Z basic-SUMO tag. Protein expression was performed in *E. coli* BL21(DE3)-Rosetta cells (Novagen). Bacteria were grown in Luria-Bertani medium at 37°C and induced with 0.3 mM isopropyl β-D-1-thiogalactopyranoside (IPTG) at 18°C overnight. Bacteria were lysed via sonication in 20 mM Tris-HCl pH 7.5, 300 mM NaCl, 1 mM PMSF, 0.1% (v/v) Triton X-100, and 20 mM imidazole. Cell lysates were centrifuged, and the supernatant was loaded onto the Ni^2+^-NTA gravity flow column (GE Healthcare) and washed in buffer 20 mM Tris-HCl pH 7.5, 300 mM NaCl and 50 mM imidazole. The target protein was washed in buffer 20 mM Tris-HCl pH 7.5, 300 mM NaCl, and 300 mM imidazole. The eluent was loaded onto a 5 mL HiTrap S HP anion-exchange column (GE Healthcare) and washed in buffer 20 mM Tris-HCl pH 7.5, 300 mM NaCl. The target protein was eluted in buffer 20 mM Tris-HCl pH 7.5, 1 M NaCl, and diluted to 20 mM Tris-HCl pH 7.5, 300 mM NaCl buffer. The N-terminal His6-Z basic-SUMO tag was removed by ULP1 protease in dialysis buffer 20 mM Tris-HCl pH 7.5, 300 mM NaCl, 5 mM β-Mercaptoethanol, and 20 mM imidazole at 4°C overnight and then removed by another 5 mL HiTrap S HP column. The target protein was further purified on a Superdex200 16/600 column (GE Healthcare) in 50 mM HEPES pH=7.5, 500 mM NaCl, 5% Glycerol, and 1 mM TCEP. The high-purity elution fractions were analyzed by SDS-PAGE and collected. The protein was flash-frozen in liquid nitrogen and stored at -80°C. The expression and purification of HCoV-OC43 NSP14, HCoV-229E NSP14, MERS-CoV NSP14, MPXV E1/E12, NSP14-10, human N7-MTase and Zika NS5 proteins were similar to that of NSP14.

### Methyltransferase (MTase) activity assay

The MTase-Glo Methyltransferase Assay Kit (Promega) was employed to detect SARS-CoV-2 NSP14 methyltransferase activity, utilizing SAM and G(5′)ppp(5′)G RNA Cap Structure Analog (New England Biolabs) as substrates. Methyltransferase activity was measured in 96-well half-area plates (Coring) via luminescence detection, utilizing a TECAN Infinite 200 PRO microplate reader for measurements.

To screen the inhibition of the MTase activity of NSP14, the assay was conducted with a reaction volume of 14 µL containing 300 nM recombinant NSP14, 1.5 µM GpppG, and SAM, and 10 µM various compounds in reaction buffer consisting of 50 mM Tris-HCl pH 8.0, 6 mM KCl, 1 mM DTT, and 1.25 mM MgCl_2_, followed by incubation at 37°C for 30 mins. 1 µL 10 × MTase Glo reagent was added to detect the production of SAH for a further incubation at 37°C for 30 mins. 5 µL MTase Glo detection buffer was added, followed by the final incubation at room temperature for 5 mins to record the luminescence. The data was analyzed by GraphPad Prism 9.

To determine the IC_50_ value of compounds, NSP14 protein was preincubated with equal volumes of compounds at final concentrations of four-fold dilution from 200 µM to 12.2 nM for 2 h on ice before conducting a total 14 µL reaction mixture that contains 300 nM recombinant NSP14 protein, 1.5 µM GpppG, and SAM, in reaction buffer consisting of 50 mM Tris-HCl pH 8.0, 6 mM KCl, 1 mM DTT, and 1.25 mM MgCl_2_, with incubation at 37 °C for 30 mins. After that, 1 µL 10 × MTase Glo reagent was added for further incubation at 37 °C for 30 mins. 5 μL of MTase Glo Detection Solution was finally added and incubated at room temperature for another 5 mins to record the reaction luminescence. Dose-dependent curves for the IC_50_ value were determined by nonlinear regression using GraphPad Prism 9.

To determine the binding mode of C10 on NSP14, the competitive enzymatic assay setup was similar, except that the SAM or GpppG concentration was increased to 7.5 µM or 37.5 µM. The inhibitory activity of the compounds against HCoV-OC43 NSP14, HCoV-229E NSP14, MERS-CoV NSP14, MPXV E1/E12, human METTL3/14, and human DNMT1 was measured using assays similar to that of NSP14.

### Thin-layer chromatography (TLC) analysis

Purified recombinant NSP14 (100 nM) was added to a 10 μL reaction mixture containing 50 mM Tris-HCl (pH 8.0), 6 mM KCl, 2 mM DTT, 1.25 mM MgCl₂, 0.1 mM SAM, and 0.1 mM GpppG, followed by incubation at 37 °C for 30 min. To assess inhibition, C10 (10 μM or 20 μM) was added to the reaction and incubated under the same conditions. As controls, 0.1 mM SAM, 0.1 mM GpppG, 0.1 mM SAH, and 0.1 mM m7GpppG (NEB) were tested in parallel. Reaction products were spotted onto polyethyleneimine (PEI) cellulose-F plates (Merck) and developed twice in 0.4 M ammonium sulfate. The migration of SAM, GpppG, SAH, and m7GpppG was visualized by UV illumination, and their relative abundance was determined by scanning the chromatogram.

### Fluorescence polarization (FP) assays

An FP-based assay using the SAM-like-FITC probe, a fluorescent analog of the methyl donor SAM, was carried out for inhibitor screening against human RNMT and Zika MTase. The reaction solution used in the assay was composed of 20 mM Tris-HCl (pH 8.0), 50 mM NaCl, 1 mM DTT, 0.05% Tween 20 and 40 nM SAM-like-FITC probe. Initially, 1 µM human RNMT or 0.5 µM Zika NS5 was incubated with DMSO or the inhibitor for 30 mins at room temperature. FP was measured by adding SAM-like-FITC probe to the reaction and using 485 nm and 535 nm for excitation and emission.

### FRET exonuclease activity

To prepare the dsRNA substrates, RNA oligos were annealed at a final concentration of 50 μM. To anneal the ssRNA oligos, samples were heated in a PCR cycler to 95 °C for 5 min and then cooled to 5 °C in 5 °C increments over 18 cycles of 1 min each. Exonuclease activity assays were performed at room temperature in black bottom 96 well plates. The reactions were performed in the following buffer: 50 mM Tris-HCl pH 7.5, 2 mM MgCl_2_, 2 mM DTT. NSP14-10 was used at 200 nM, while the dsRNA substrate was added at a final concentration of 100 nM. The fluorescence intensity of each well was measured every 60 s on an Infinite 200 PRO microplate reader (Tecan) over the course of the activity assay using the following settings: Excitation 490 nm (±9 nm)/ Emission 520 nm (±20 nm). Top Cy3 RNA (5’-3’): Cy3-GGUAGUAAUCCGCUC; Bottom quencher RNA (5’-3’): UUUUUUUUUUUUUUUUUUUUGAGCGGAUUACUACUACC-BHQ-2.

### Surface plasmon resonance (SPR) binding assays

The Biacore 8K instrument (GE Healthcare) was employed to determine the binding kinetics of compounds at 25 °C. The protein was immobilized on a Series S CM5 sensor chip by a standard coupling procedure. Compounds were diluted serially and injected for 60 s (contact phase), followed by 120 s (dissociation phase) at a flow rate of 30 μL/min. The binding kinetics rate constants, K_a_ (M⁻¹ s⁻¹), K_b_ (s⁻¹), and the equilibrium dissociation constant K_D_ (M) were calculated using the Biacore 8K Evaluation software (1.1). To determine the equilibrium dissociation constant K_D_, response data at equilibrium binding were processed by the software mentioned above. Response data at equilibrium binding were also processed to determine the equilibrium dissociation constant K_D_. The competitive binding assay setup was similar, except that 20 µM SAM, GpppG, SAH and m7GpppG was added to the running buffer. The final curves were visualized using GraphPad Prism 9.

### Differential scanning fluorimetry (DSF) assay

DSF assays were performed using the CFX Connect Real-Time System (Bio-Rad). Each 25 µL reaction solution contained 10 µM NSP14 protein in the buffer (20 mM Tris-HCl pH 7.5, 300 mM NaCl, and 1 mM DTT), 25× SYPRO (Supelco, 5000× concentrated solution in DMSO, S5692) orange, and tested concentrations of compounds (5% v/v DMSO), and was heated from 25°C to 85°C. Fluorescence intensities were recorded every 0.5°C / 30 s. The data from the Bio-Rad tests were analyzed and then fitted with the Boltzmann equation in GraphPad software to calculate the melting temperature (*T_m_*).

### Simulation system construction

The crystal structure of NSP14 from SARS-CoV-2 in complex with SAH was retrieved from RCSB protein data bank with PDB entry of 7R2V. To build the ternary complex of NSP14-m7GpppG-SAH for ligand binding mode exploration, crystal structure of NSP14-m7GpppG complex with PDB entry of 7QIF was utilized to determine the binding site of m7GpppG via structural superimposition. Schrodinger2020 package was used for the simulation system construction. Herein, the ternary complex was processed by Protein Preparation Wizard module, including hydrogen atoms adding, bond orders assigning, hydrogen bond optimizing. To improve the structural reliability, the prepared structure was minimized under OPLS3e force filed with the root-mean-square deviation (RMSD) of the heavy atoms converged to 0.3 Å. The chemical structure of C10 was processed by LigPrep module, and the docking grid was generated by Receptor Grid Generation module referring to SAH molecule. Subsequently, the binding mode of C10 was investigated via molecular docking using Glide module, and the docking complex with suitable binding mode as well as favorable docking score was selected for further exploration.

### Molecular dynamics simulations

The constructed ternary complex of NSP14-m7GpppG-SAH and the selected docking complex of C10 were projected to conventional MD simulations with Amber24 package. The AMBER ff19SB force filed was applied to the protein, and the AMBER general force field 2 (GAFF2) was applied to the ligands with the AM1-BCC charges. The complexes were solvated into a TIP3P water cubic box, and counter ions were added to neutralize the net charge of the systems. The Particle Mesh Ewald (PME) algorithm was employed for the long-range electrostatic interaction, and the cutoff for the long-range electrostatic interaction, and the cutoff for the real-space interactions was set to 12 Å. Subsequently, each system was subjected to a stepwise structure minimization. Firstly, the water and counter ions were optimized by a 5000 steps of steepest descent and 5000 steps of conjugated gradient minimization with other atoms restrained by 50 kcal·mol^-1^·Å^-2^. Secondly, the constraint potential was decreased to 10 kcal·mol^-1^·Å^-2^ for another 5000 steps of steepest descent and conjugated gradient minimization. Thirdly, the system underwent another 5000 steps of steepest descent and conjugated gradient minimization, wherein only the protein atoms were restrained by 10 kcal·mol-1·Å^-2^ potential. Finally, with all restraint potential removed, the system was further optimized by 5000 steps of steepest descent and conjugated gradient minimization. Then, with 5 kcal·mol-1·Å^-2^ potential applied to protein backbone atoms, the system was gradually heated to 300 K over a period of 100 ps under the NVT ensemble, which was further equilibrated for 100 ps in the NPT ensemble with protein and ligand restrained by 1 kcal·mol-1·Å^-2^, respectively. Finally, three replicas of unrestrained 200 ns production simulation were performed in the NPT ensemble using the pmemd.cuda module, and the simulation trajectory were saved every 10 ps. The root-mean-square deviation (RMSD) analyses were conducted by the cpptraj module in AmberTools 24. Moreover, to reveal the binding pattern of C10, the protein-ligand binding free energy was calculated with the MM/GBSA free energy decomposition via the MMPBSA.py script using the final 20 ns equilibrated simulation trajectory.

### Hydrogen-deuterium exchange mass spectrometry (HDX-MS)

The HDX-MS experiment was performed using a Tris-Hcl buffer at pH 7.4, with two parallel buffers prepared using either water or deuterium oxide as the solvent. Sample preparation was automated and efficiently carried out using the PAL3 System (LEAP Technologies).

For continuous labeling, 4 μL of the NSP14 (12 μM), either in the presence or absence of C10, was mixed with 30 μL of D_2_O buffer. The mixture was incubated at 10 °C for 10, 60, and 600 seconds to allow hydrogen-deuterium exchange under controlled conditions. The deuterium-exchanged samples were quenched with 30 µL of ice-cold quench buffer (4 M guanidine hydrochloride, 200 mM TCEP, 100 mM citric acid, pH 2.3). Fifty microliters of quenched samples were thawed and immediately injected onto an immobilized pepsin column (2.1 × 30 mm; Thermo Fisher Scientific, Waltham, MA, USA) at a flow rate of 50 µL/min with 0.1% formic acid in H_2_O at 4 °C. Peptide fragments were collected on an Acclaim PepMap300 C18 column (5 µm, 1.0 mm × 15 mm; Thermo Fisher Scientific) for desalting with 0.1% formic acid in H_2_O, followed by isolation via liquid chromatography using an ACQUITY UPLC Peptide CSH C18 column (130 Å, 1.7 µm, 1 mm × 50 mm; Waters, Milford, MA, USA). Chromatographic separation was performed at a flow rate of 45 µL/min with an acetonitrile gradient starting at 5% and increasing to 80% over 10 minutes. To minimize deuterium back-exchange, the system, including the trapping and UPLC columns, was maintained at 0.5 °C, and all buffers were adjusted to pH 2.5.

Mass spectral analyses were carried out using an Orbitrap Fusion Tribrid™ Mass Spectrometer (Thermo Fisher Scientific, USA) equipped with a heated electrospray ionization (HESI) source in positive ion mode. Tandem MS (MS/MS) data were processed using BioPharma Finder 2.0 software (Thermo Fisher Scientific, USA) for peptide identification, and HDX-MS data were analyzed with HDExaminer 3.0 software (Sierra Analytics, Modesto, CA, USA). Per-residue deuterium uptake differences were calculated for each time point, manually validated, and exported based on overlapping peptides. Statistical significance for differential HDX data was assessed using an unpaired t-test for each time point. All data were obtained from at least three independent experiments.

### Cell culture

To construct the lentivector expressing the human ACE2 gene, the human ACE2 gene (Miaolingbio #P5271) was PCR-amplified and cloned into the pLVX-EF1α-IRES-blast (Addgene #85133). A549 (ATCC #CCL-185) cells were cultured at 37°C in Dulbecco’s modified Eagle’s medium (Hyclone #SH30243.01) supplemented with 10% fetal bovine serum (FBS), 10 mM HEPES, 1 mM sodium pyruvate, 1× non-essential amino acids, and 100 U/mL of penicillin-streptomycin. The A549-ACE2 cell line was generated by transduction of the lentivector expressing the human ACE2 gene.

HEK293T (ATCC #CRL-3216), HeLa (ATCC #CCL-2), and BHK-21 (ATCC #CCL-10) and A549 (ATCC #CCL-185) were cultured at 37°C in Dulbecco’s modified Eagle’s medium supplemented with 10% fetal bovine serum. The plasmid pLVX-EF1α-HA-SARS-CoV-2 NSP14-IRES-EGFP-Puro was constructed and sequenced. The day before transfection, 1×10⁶ cells per well were plated in 6-well plates. Then, the cells were transfected with PEI (PolySciences, cat#23966-1). Briefly, 2000 ng of recombinant DNA was incubated at room temperature for 20 minutes with PEI, with PEI:DNA = 3:1 in 100 µL serum-free medium, and then the mixture was added to the cells. After transient transfection for 8 hours, the medium was removed and replaced with medium supplemented with 10% fetal bovine serum.

### Cellular thermal shift assay (CETSA)

The recombinant plasmid pLVX-EF1α-HA-NSP14-IRES-EGFP-Puro was transiently transfected into HEK293T (ATCC #CRL-3216) mammalian cells when the cell density reached 60%. The transfection was performed in serum-free DMEM medium using a plasmid concentration of 1 µg/mL (plasmid:PEI = 1:3). After 6 hours of transfection, the medium was replaced with DMEM containing 10% serum. After 40 hours, 20 µM of C10 was added, and the cells were incubated for an additional 8 hours before harvesting. The cells were then washed and resuspended in PBS. Afterwards, treated cells were equally divided and incubated at various temperatures for 3 mins, then cooled at room temperature for 3 mins. Samples were lysed by repeated three cycles of liquid nitrogen flash freezing and thawing. Cell lysates were centrifuged at 17,500 rpm at 4°C for 15 minutes, and the supernatants were analyzed by western blot. The concentration-dependent CETSA assay was similar, except that the cells were incubated with various concentrations of C10 or DMSO and incubated at 54°C for 3 mins.

### Time-of-addition experiment

The C10 (3 µM, 10 µM and 20 µM) was used for the time-of-addition experiment. A549-ACE2 cells were treated with C10 or DMSO at different stages of SARS-CoV-2 trVLP infection. For “full-time” treatment, A549-ACE2 cells were pre-treated with the C10 for 1h prior to virus infection, followed by incubation with virus for 2h in the presence of C10. Then, the virus-C10 mixture was removed. Cells were washed with PBS, and further cultured with drug-containing medium until the end of the experiment. For “entry” treatment, C10 was added to the cells for 1h before virus infection and maintained during the 2h viral attachment process. Then, the virus-C10 mixture was replaced with fresh culture medium without C10 till the end of the experiment. For “post-entry” experiment, virus was added to the cells to allow infection for 2h, and then virus-containing supernatant was replaced with C10-containing medium until the end of the experiment. The experimental condition of the DMSO-treatment group was consistent with that of the “full-time” group. For all the experimental groups, cells were infected with virus at the MOI of 0.02, and at 24 h.p.i., the luciferase activity was measured using the Nano-Glo Luciferase Assay kit (Promega 1040#N1110).

### Selection for SARS-CoV-2 resistance mutations

To select for C10 resistance mutations in SARS-CoV-2 trVLP was passaged on Vero-N cells in the presence of DMSO (control lineage) or increasing concentrations of C10. In detail, Vero-N cells were seeded in 12-well plate in complete medium (DMEM supplemented to contain 10% FBS). The next day, cells were exposed to SARS-CoV-2 trVLP at an MOI of 1 PFU at 37 °C for 2 h. Virus was removed, and complete medium containing either DMSO or 10 μM C10 was added to cells. Cells were incubated at 37 °C for 72 h. Supernatants were collected and used to inoculate fresh Vero-N cells. Virus was passaged serially in this manner for independent replicates, increasing the concentration of C10 until it reached 20 μM. After 5 passages, virus isolates recovered from each lineage were then characterized for their resistance to 10 μM or 20 μM C10 by determining the fluorescence.

To identify C10-resistance mutations emerging in SARS-CoV-2 trVLP passaged over 5 passages in Vero-N cells, total RNA was isolated from cell culture supernatants from each viral lineages using TRIzol Reagent according to the manufacturer’s instructions. RNA was further purified using the RNA simple Total RNA Extraction Kit (Tiangen, DP419). cDNA was prepared using a transcription kit (TransGen Biotech). For PCR amplification of the NSP14 coding region the following primers were used: NSP14-forward (5’-GAAAATGTAACAGGACTCTTTAAAGATTGTAGTAAGG-3’) and NSP14-reverse (5’-CTGAAGTCTTGTAAAAGTGTTCCAGAGG-3’). PCR reactions were purified using the PCR product extraction magnetic beads (Biolinkedin, L-3001) according to manufacturer’s instructions, and linear amplicon sequencing was performed by NGS.

### Competitive fitness assay

Vero-N cells were infected with purified SARS-CoV-2 trVLP (wild-type or NSP14-V290A) at 37 °C for 2 h. After adsorption, the inoculum was removed, and cells were washed three times with 1× PBS to eliminate residual virus, followed by the addition of complete medium. Supernatants were collected and used for subsequent passages. Viral RNA from the input inoculum (P0) and each passaged supernatant (P1 and P2) was extracted using TRIzol Reagent (Invitrogen) according to the manufacturer’s instructions. Purified RNA was reverse transcribed to cDNA, and the NSP14 coding region was amplified by PCR. Amplicons were subjected to next-generation sequencing (NGS) for mutation analysis.

### Next-Generation Sequencing

For each sample, more than 2µg purified PCR fragments (1kb∼10kb) was used for library preparation. Fragmented DNAs were treated with End Prep Enzyme Mix for end repairing, 5’Phosphorylation and 3’dA-tailing in one reaction, followed by a T-A ligation to add universal hairpin adapters to both ends. Then fragment size of the libraries was analyzed by Agilent Bioanalyzer (Agilent Technologies, Palo Alto, CA, USA), and quantified by Qubit 3.0 Fluorometer. The final SMRTbell libraries were loaded on PacBio Sequel IIe instrument according to manufacturer’s instructions (Pacific Biosciences of California, Inc., California, USA). Perform statistical analysis on hifi reads data. Split CCS reads according to barcode by using lima software (version 1.9.0), and statistics the split results. If the target area is specified, the target sequence is captured according to the 10nt of upstream and downstream of the target sequence, otherwise, the target sequence refs to the full-length sequences. The CCS reads are aligned to the reference sequence by using BWA software, and the section on the alignment is extracted and mutation abundance information is counted. The target sequence is translated into an amino acid sequence starting from the first base, and abundance statistics are performed. Extract amino acid sequences of specified lengths and record the amino acid distribution at each position. Mutations were detected using the sentieon software based on the comparison results of the bwa software.

### RNA-seq assay

A total amount of 2 μg RNA per sample was used as input material for the RNA sample preparations. After 48 hours of treatment with compounds or DMSO on HEK293T cells expressing NSP14. Sequencing libraries were generated using the NEBNext Ultra RNA Library Prep Kit for Illumina (#E7530L, NEB, USA) following the manufacturer’s recommendations, and index codes were added to attribute sequences to each sample. Briefly, mRNA was purified from total RNA using poly-T oligo-attached magnetic beads. Fragmentation was carried out using divalent cations under elevated temperature in the NEBNext First Strand Synthesis Reaction Buffer (5×). The library fragments were purified using QiaQuick PCR kits and eluted with EB buffer. The aimed products were retrieved, PCR was performed, and the library was completed. Subsequently, the RNA concentration of the library was measured using the Qubit® RNA Assay Kit on the Qubit 3.0 to preliminarily quantify it, and then diluted to 1 ng/μL. IInsert size was assessed using the Agilent Bioanalyzer 2100 system (Agilent Technologies, CA, USA), and the qualified insert size was accurately quantified using the StepOnePlus Real-Time PCR System (library valid concentration > 10 nM). Finally, clustering of the index-coded samples was performed on a cBot cluster generation system using the HiSeq PE Cluster Kit v4-cBot-HS (Illumina) according to the manufacturer’s instructions. After cluster generation, the libraries were sequenced on an Illumina platform, and 150 bp paired-end reads were generated.

### Quantitative Real-Time PCR (qPCR)

After 40 hours of C10 or DMSO treatment of HEK293T-NSP14 cells or SARS-CoV-2 trVLP infected A549-ACE2 cells, total RNA was extracted from the cells using the Trizol reagent (Invitrogen), and cDNA was transcribed using a transcription kit (TransGen Biotech). qRT-PCR analysis was performed using the SYBR Green (Bio-Rad) method on the ABI Fast 7500 Real-Time PCR instrument (PerkinElmer Applied Biosystems). Gene expression of AURKA1 (primers from Beyotime Biotechnology) was calculated using the comparative ΔCt method with GAPDH (primers from Beyotime Biotechnology) for normalization. Gene expression of Spike (SARS-CoV-2-Spike-F: CCTACTAAATTAAATGATCTCTGCTTTACT, SARS-CoV-2-Spike-R: CAAGCTATAACGCAGCCTGTA) was calculated using the comparative ΔCt method with HRPT1 (primers from Beyotime Biotechnology) for normalization. Gene expression of RSAD2, IFI6, IRF7, IFIT2 and OAS1 (primers from Beyotime Biotechnology) were calculated using the comparative ΔCt method with RPS11 (primers from Beyotime Biotechnology) for normalization. Gene expression of IMPDH2 (IMPDH2-F: AGTGGCTCCATCTGCATTACGC, IMPDH2-R: GGATTCCTCCATCAGCAATGACC) was calculated using the comparative ΔCt method with GAPDH (primers from Beyotime Biotechnology) for normalization.

### Polysome profiling experiment

A549-ACE2 cells were seeded in T75 flasks at a density of 5 × 10^6^ cells per flask. The following day, cells were mock-infected or infected with SARS-CoV-2 trVLP at an MOI of 1 PFU/cell. After 2 h of incubation, the inoculum was removed, cells were washed with 1× PBS, and fresh medium containing DMSO, 20 μM C10, or 40 μM C10 was added. At 8 hours post-infection (hpi), cycloheximide (100 μg/mL in RNase-free ethanol) was added for 5 min to arrest translation, after which cells were lysed in polysome lysis buffer [10 mM Tris-HCl (pH 8.0), 150 mM NaCl, 15 mM MgCl₂, 20 mM DTT, 1% Triton X-100, 150 μg/mL cycloheximide] supplemented with RNasin (Promega).

Lysates were clarified by centrifugation (12,000 × g, 10 min, 4 °C), and total RNA concentrations were determined by NanoDrop 2000 (Thermo Fisher Scientific). Equal amounts of RNA (200 μg) were diluted to 300 μL and layered onto 12 mL linear sucrose gradients (10–50%). Gradients were centrifuged at 38,500 × g for 2.5 h at 4 °C in a Beckman SW41Ti rotor using an Optima XPN-100 ultracentrifuge. Fractions (∼0.65 mL each) were collected at 45 s intervals using a piston gradient fractionator (Bio-Comb) with continuous monitoring of absorbance at 260 nm. RNA from free and polysomal fractions was extracted with TRIzol, reverse-transcribed (TransGen Biotech), and analyzed by qRT-PCR using SYBR Green (Bio-Rad) on an ABI 7500 Fast Real-Time PCR system. SARS-CoV-2 Spike mRNA levels were measured in technical triplicates and normalized to HPRT1 mRNA using the ΔCT method, with DMSO-treated samples as reference.

### Western blot

For CETSA samples, an equal volume of reducible loading buffer (50 mM Tris, pH 6.8, 10% Glycerol, 2% SDS, 0.05% [wt/vol] Bromophenol Blue, 100 mM DTT) was added to each 100 μL of supernatants. For transcriptome samples, after 48 hours of treatment with compounds or DMSO on HEK293T cells expressing NSP14, 100 μL of lysate (RIPA: PMSF: DNAse: protease inhibitor: phosphatase inhibitor = 100:1:1:1:1) was added, shaken on ice for 1 hour, and then centrifuged at 14,000 rpm for 10 mins. Supernatants were quantified using a BCA assay, diluted with PBS to a 40 μL system, and mixed with an equal volume of loading buffer.

An equal volume of samples was loaded for electrophoretic separation, followed by wet-transfer to the PVDF film. The PVDF film was blocked at room temperature for 1 hour with 5% skim milk in TBST (100 mM NaCl, 10 mM Tris, pH 7.6, 0.1% Tween 20), and then incubated with the corresponding primary antibodies at 4 °C overnight. The PVDF films were washed with TBST and incubated at room temperature for 1 hour with horseradish peroxidase (HRP)-labeled secondary antibodies, followed by additional TBST washes. Protein strips were exposed using SuperPico ECL Master Mix (Vazyme) according to the manufacturer’s instructions with the Amersham ImageQuant 800 (Cytiva). Data were processed using ImageJ and labeled with Adobe Illustrator.

(anti-HA: CST3724 1:1000; GAPDH: Santa Cruz sc-32233 1:10,000; AURKA: Proteintech 82906-1-RR 1:1000; CDK1: Proteintech 19532-1-AP 1:2000; Cyclin B1: Proteintech 67686-1-Ig 1:2000; Beta-Actin: Proteintech HRP-66009 1:10,000; Anti-SARS-CoV-2 Spike glycoprotein S1 (RBD): GB12869-50 1:1000; Peroxidase AffiniPure Goat Anti-Mouse IgG: Jackson Immuno Research, 115-035-003 1:5000; Peroxidase AffiniPure Goat Anti-Rabbit IgG: Jackson Immuno Research, 111-035-003 1:5000).

### Cell cycle analysis by flow cytometry

Cells were fixed in 70% ethanol overnight at 4°C, then washed with PBS and stained with PI/RNase Staining Buffer Solution (Biosharp) for 30 mins in the dark. The cell cycle distribution was analyzed using a CytoFLEX LX flow cytometer. Data analysis was carried out using ModFit LT 5.0.

### Cell viability assay

A CellTiter-Lumi Steady Cell Viability Assay (Beyotime # C0069M) was conducted according to the manufacturer’s instructions. Approximately 1 × 10^4^ A549-ACE2 cells per well were plated in 96-well plates one day before the test. Compounds were three-fold diluted to a final concentration ranging from 200 μM to 10.16 nM in DMEM medium containing 2% FBS. The diluted compounds were added to the cells and cultured for 24 hours. Subsequently, after removing the supernatants, the CellTiter-Lumi reagents were added to each well, followed by shaking at room temperature for 2 mins and incubation in the dark for 10 mins. Luminescence was measured using a FlexStation 3 (Molecular Devices) with an integration time of 0.5 seconds per well.

### Viral infection assay and immunofluorescence staining

A549-ACE2 cells were plated in 96-well plates one day before the experiment. Diluted compounds were added to the cells as pretreatment for 1 hour, then SARS-CoV-2 original (MOI 0.2), Delta (MOI 0.5), and Omicron (MOI 0.1) strains were used to infect A549-ACE2 cells for 24 hours, while HCoV-OC43 and HCoV-229E strains were used to infect A549-ACE2 cells with an MOI of 0.2 for 48 hours. Subsequently, cells were fixed with 4% PFA in PBS for 30 minutes, and permeabilized with 0.2% Triton X-100 for 1 hour. Cells were then incubated with house-made mouse nucleocapsid protein serum (1:1000) against different coronaviruses at 4 °C overnight. After three washes, cells were incubated with the secondary goat anti-mouse antibody conjugated with Alexa Fluor 555 (Thermo #A-21424, 2 μg/mL) for 2 hours at room temperature, followed by staining with 4′,6-diamidino-2-phenylindole. Images were collected using an Operetta High Content Imaging System (PerkinElmer) and processed using the PerkinElmer Harmony high-content analysis software v4.9 and ImageJ v2.0.0. All experiments involving SARS-CoV-2 virus infection were performed in the biosafety level 3 (BSL-3) facility of Fudan University following the approved standard operating procedures.

### SARS-CoV-2 replicon assay

The replicon system was constructed as previously described [2]. Briefly, the full-length viral RNA of SARS-CoV-2 strain SH01 was reverse-transcribed and PCR-amplified into 6 fragments. The region encompassing the spike gene to ORF8 was replaced with a Nanoluciferase-P2A-puromycin cassette, and the T7 promoter was inserted upstream of the viral genome to initiate transcription. All fragments were assembled into the pBeloBAC11 vector in vitro as one plasmid and then amplified in bacteria. Replicon RNA was transcribed from the single plasmid using the mMESSAGE mMACHINE T7 Transcription Kit (Invitrogen #AM1344) according to the manufacturer’s instructions. RNA was then electroporated into HeLa cells using the Invitrogen Neon NxT Electroporation System to assess the virus replication efficiency. After electroporation, the cells were seeded in a 96-well plate and immediately treated with 10 μM C10, DMSO as a negative control, and remdesivir (10 μM) as a positive control. At 24 hours post-transfection, the luciferase activity was measured using the Nano-Glo Luciferase Assay kit (Promega #N1110).

### Pharmacokinetic analysis in vivo

Male ICR mice (6–8 weeks, 18–20 g) obtained from Hangzhou Hangsi Biotechnology Limited Company were used to evaluate the pharmacokinetic profile of various compounds. Compounds were dissolved in 5% dimethyl sulfoxide and 95% cyclodextrin solution (m(cyclodextrin):m(normal saline) = 1:2). After oral administration and intraperitoneal injection of compounds, blood samples were collected into EDTA-containing tubes by retro-orbital bleeding at 0.25, 0.5, 1, 2, 4, 8, 12, and 24 hours. Plasma was obtained by centrifugation at 4000 rpm for 10 mins. The concentrations of various compounds were quantified by LC-MS. Compound concentrations were determined with an API 4000 + LC-MS system. These animals were bred at the College of Pharmaceutical Sciences, Zhejiang University. All animal studies were performed in strict accordance with the institutional guidelines as defined by the Institutional Animal Care and Use Committee, Zhejiang University Laboratory Animal Center (Hangzhou, China).

### Toxicity study

According to the pharmacokinetic characteristics of C10, intraperitoneal injection (I.P.) administration to C57BL/6 mice was designed in the absence of virus. Doses of 300 mg/kg, 200 mg/kg, and 100 mg/kg were continuously administered twice a day (B.I.D.) for three days, followed by recording changes in body weight and mouse survival.

### Animal experiments

Six- to eight-week-old male K18-hACE2 transgenic mice were used in the study in the BSL-3 laboratory at Fudan University. The experimental protocol has been approved by the Animal Ethics Committee of the School of Basic Medical Sciences at Fudan University. The mice were inoculated intranasally with 5 × 10³ focus-forming units of SARS-CoV-2 nCoV-SH01 strain (GenBank accession no. MT121215). Animals were briefly anesthetized for the intranasal inoculation using inhaled isoflurane. Animals were monitored throughout the inoculation and anesthetic recovery periods.

The compound (C10), vehicles, or remdesivir were administered via intraperitoneal injection. Twice-daily treatment for all groups began 1 hour post infection on day 1 and continued through day 3. Mice were euthanized on day 4. Lungs were collected on day 4 following euthanasia. Tissues were weighed, homogenized in DMEM, and centrifuged at 3500–4000 rpm for 5 mins. Cleared supernatants were collected for focus-forming. Briefly, Vero E6 monolayer in 96-well plates was inoculated with serially diluted virus for 2 hours and then overlaid with methyl cellulose for 48 hours. Cells were fixed with 4% PFA in PBS for 1 h and permeabilized with 0.2% Triton X-100 for 1 h. Cells were stained with homemade mouse serum against the N protein of SARS-CoV-2 overnight at 4 °C, incubated with the secondary goat anti-rabbit HRP-conjugated antibody (Thermo Fisher #31460) for 2 h at room temperature. The focus-forming units were developed using TrueBlue substrate (Sera Care #5510-0030). All experiments involving SARS-CoV-2 virus infection were performed in the biosafety level 3 (BSL-3) facility of Fudan University following the approved standard operating procedures.

### Lung Histology and Immunohistochemistry (IHC)

Lung tissues were collected and fixed in 4% paraformaldehyde (PFA) for 48 hours. Following fixation, tissues were embedded in paraffin, sectioned, and stained with hematoxylin and eosin (H&E) for morphological assessment. For viral antigen detection, sections were incubated with an in-house mouse anti-SARS-CoV-2 nucleocapsid protein serum (1:1000) followed by an HRP-conjugated goat anti-mouse IgG secondary antibody (1:5000 dilution, Invitrogen). Microscopic analysis was performed using an Olympus microscope (Tokyo, Japan). Histopathological evaluation was conducted by a board-certified veterinary pathologist, assessing two parameters: congestion and alveolar wall thickening. Both parameters were scored using an ordinal scale based on the percentage of affected tissue: 0 (none), 1 (<25%), 2 (26-50%), 3 (51-75%), and 4 (>75%). Similarly, IHC staining was evaluated based on the percentage distribution of antigen staining in lung tissues. All experiments involving SARS-CoV-2 virus infection were performed in the biosafety level 3 (BSL-3) facility of Fudan University following the approved standard operating procedures.

### The biosafety level (BSL) used for experiments

All experiments involving SARS-CoV-2 trVLP virus infection were performed in the biosafety level 2 (BSL-2) facility of Fudan University. All experiments involving SARS-CoV-2 virus infection were performed in the biosafety level 3 (BSL-3) facility of Fudan University following the approved SOPs. SARS-CoV-2 replicon assay was performed in the biosafety level 2 (BSL-2) facility of Fudan University.

**Chemistry:**

**in Source data.**

---

**Extended Data Figure 1.**
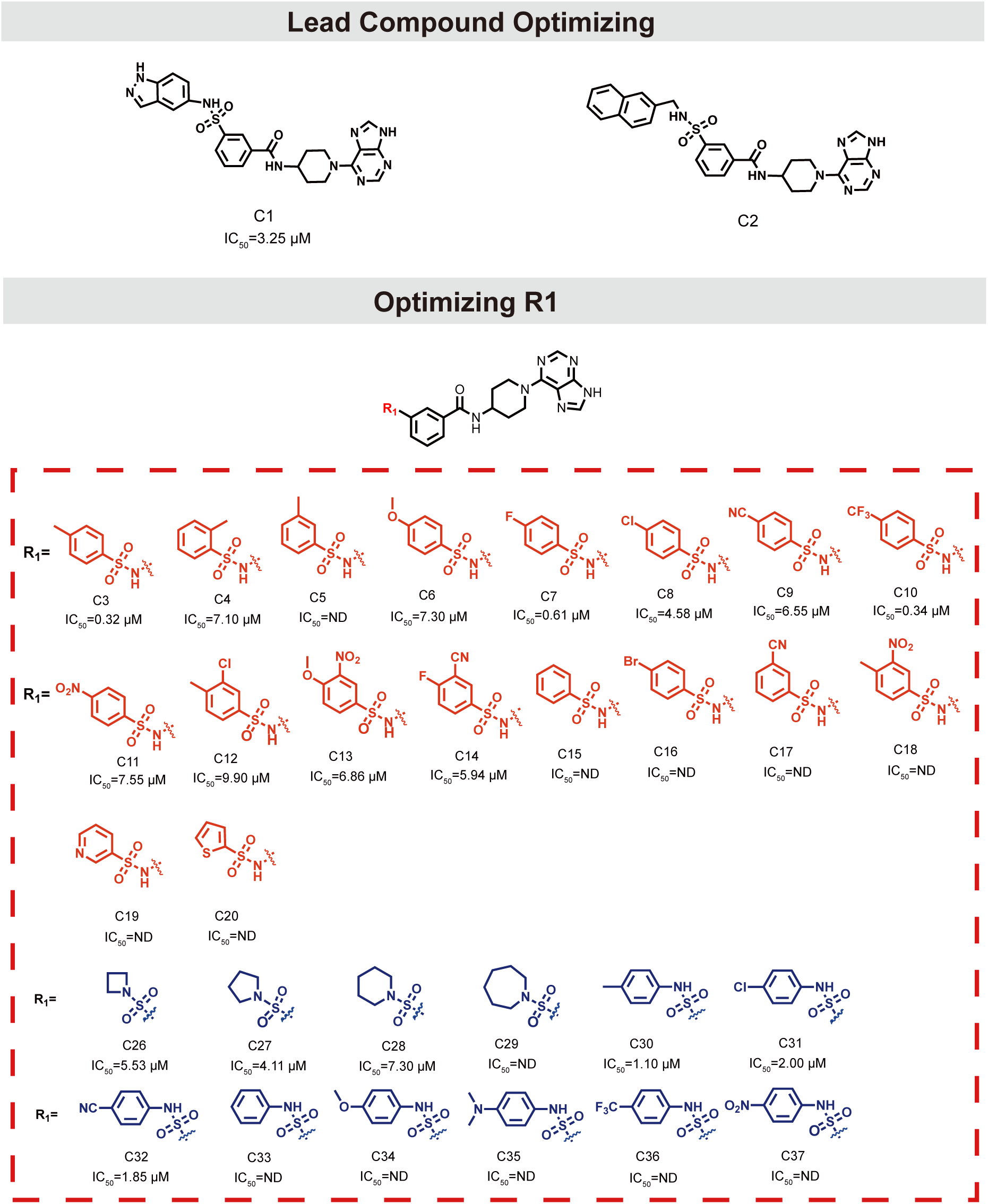
Stepwise structural optimization procedure for C chemotype compounds.

**Extended Data Figure 2.**
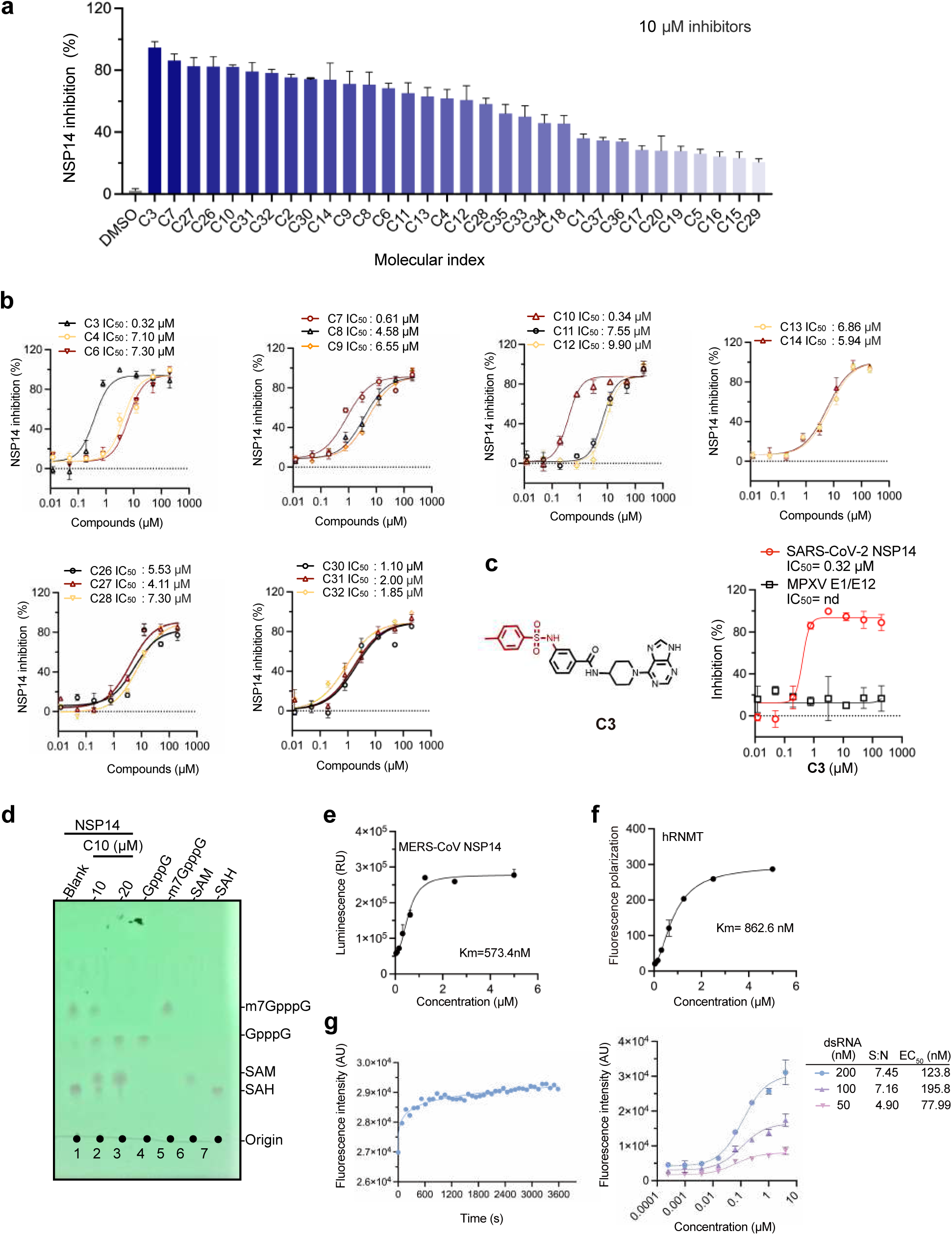
Enzymatic inhibition of C chemotype compounds. **a.** Histogram showing the inhibition of SARS-CoV-2 NSP14 by C-family compounds at 10 μM, as measured by the Mtase-Glo assay. Data are presented as mean ± SD from three independent replicates. **b.** Dose-dependent inhibition curves of top-performing C chemotype compounds against SARS-CoV-2 NSP14, as measured by the Mtase-Glo assay. **c.** Chemical structure of hit compound C3 (Left). Dose-dependent inhibition curves of C3 against SARS-CoV-2 NSP14 and MPXV E1/E12 as measured by the Mtase-Glo assay (Right). Data are presented as mean ± SD from three independent replicates. **d.** TLC analysis of m7GpppG generated from GpppG and SAH generated from SAM after treatment with NSP14 (lanes 1–3). Addition of 10 μM or 20 μM C10 inhibited NSP14 activity, as shown in lanes 2–3. The migration positions of SAM, SAH, GpppG, and m7GpppG are indicated on the right. **e.** Titration of MERS-CoV NSP14 at 1.5 µM SAM and 1.5 µM GpppG for Km value determination. N = 3. Data are shown as mean ± SD. **f.** Titration of hRNMT for Km value determination as measured by the FP assay. N = 3. Data are shown as mean ± SD. **g.** Time-dependent NSP14-NSP10 exonuclease activity curves measured by the FRET assay (left). NSP14-NSP10 concentration-dependent exonuclease activity curves with 50 nM, 100 nM, and 200 nM dsRNA substrate (right).

**Extended Data Figure 3.**
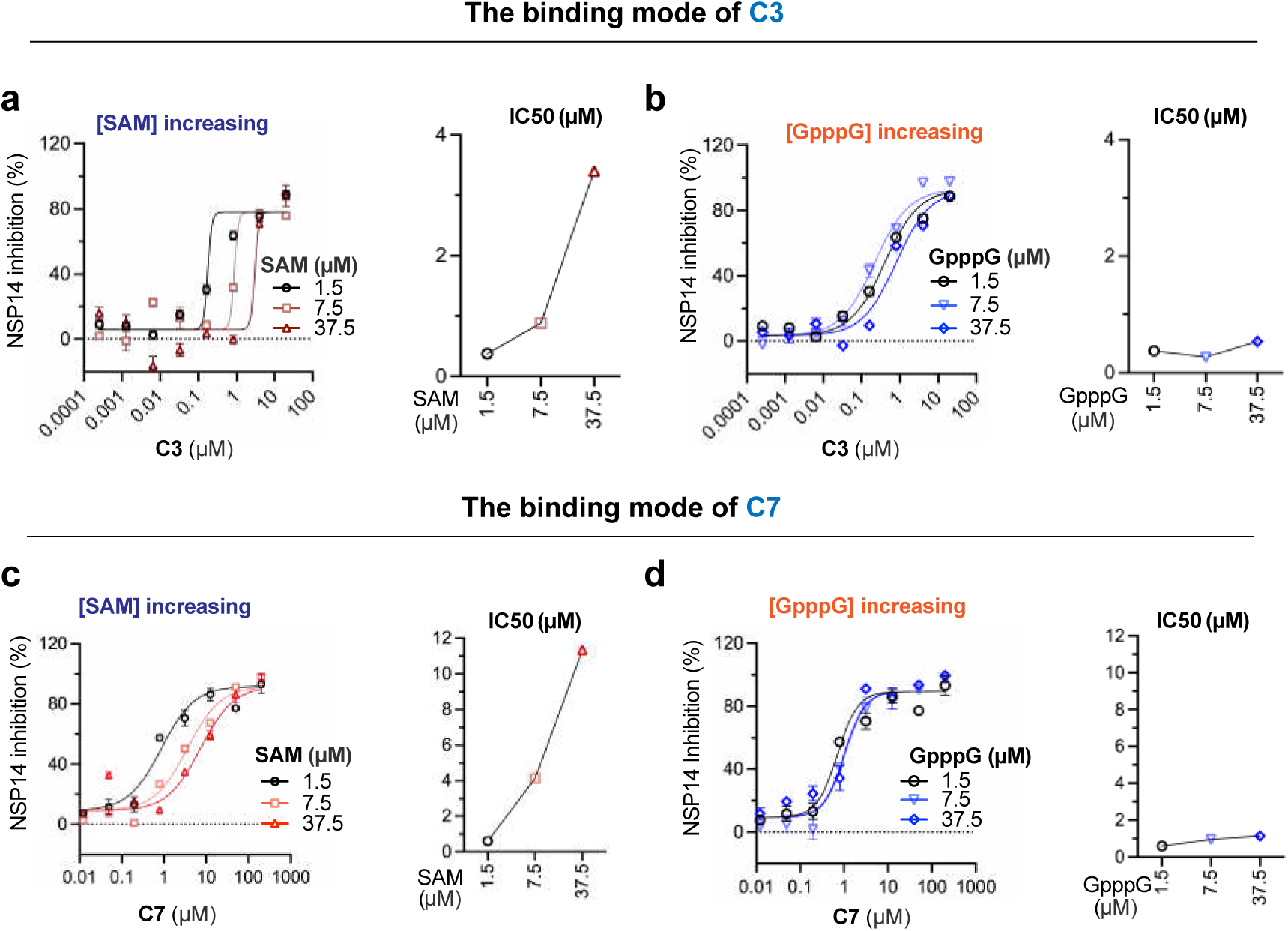
Inhibitory mode of C3 and C7 compounds against SARS-CoV-2 NSP14. **a-d.** Dose-dependent inhibition curves of C3 (a, b) and C7 (c, d) against NSP14 with varying concentrations of GpppG and SAM. In a and c, SAM concentrations were increased while GpppG concentration was fixed at 1.5 μM. In b and d, GpppG concentrations were increased while SAM concentration was fixed at 1.5 μM. The corresponding IC_50_ values are presented as scatter plots. Data are shown as mean ± SD from three independent replicates.

**Extended Data Figure 4.**
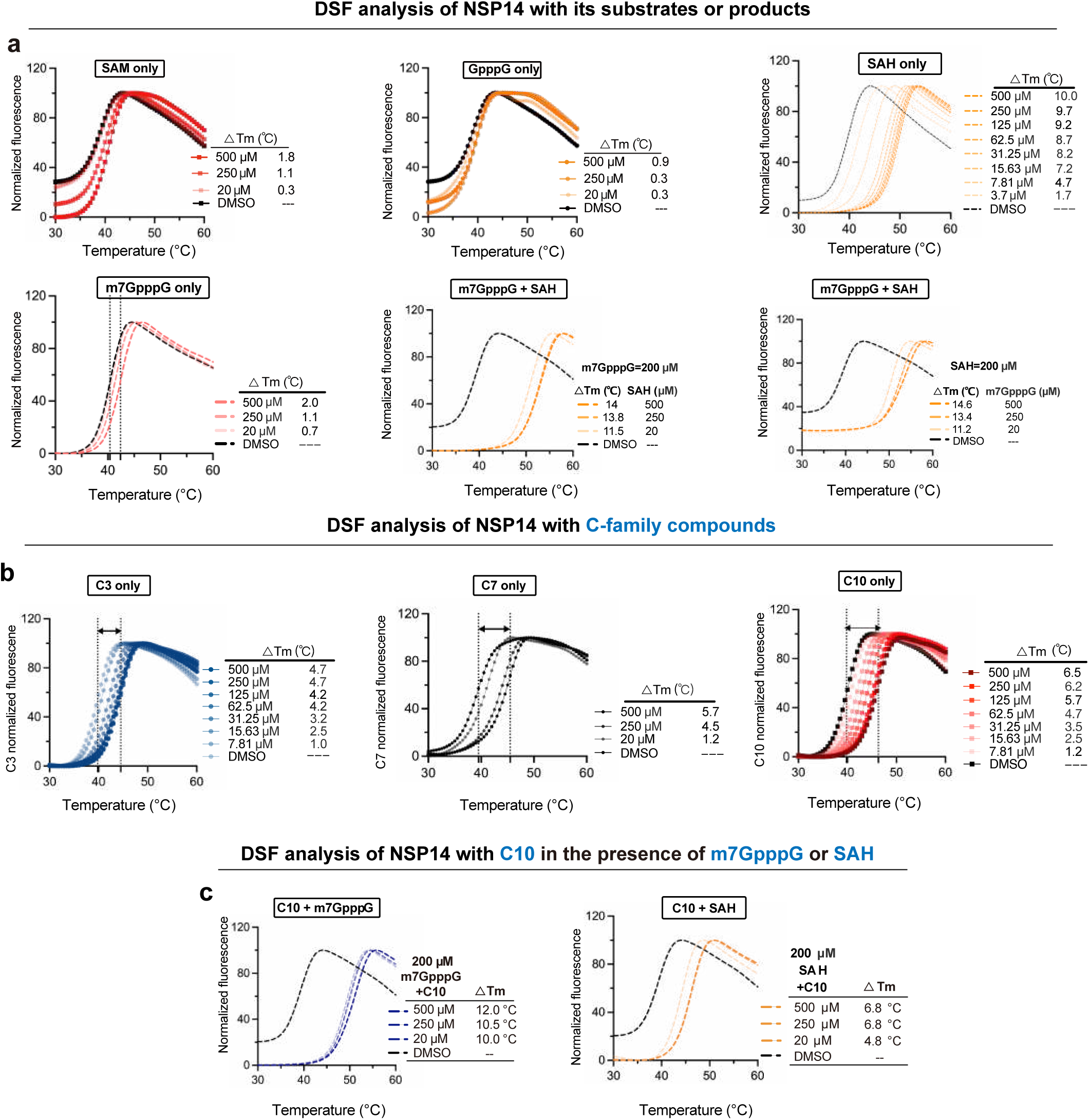
Melting temperature shifts of NSP14 measured by DSF assays. **a.** DSF analysis of NSP14 in the presence of its substrates (SAM and GpppG), products (SAH and m7GpppG), and the product combination (SAH + m7GpppG). **b.** DSF analysis of NSP14 with C-family compounds (C3, C7, and C10) **c.** In the presence of 200 μM m7GpppG, instead of 200 μM SAH, treatment with C10 caused an additional increase in the Tm of NSP14, exhibiting a synergistic effect between C10 and m7GpppG.

**Extended Data Figure 5.**
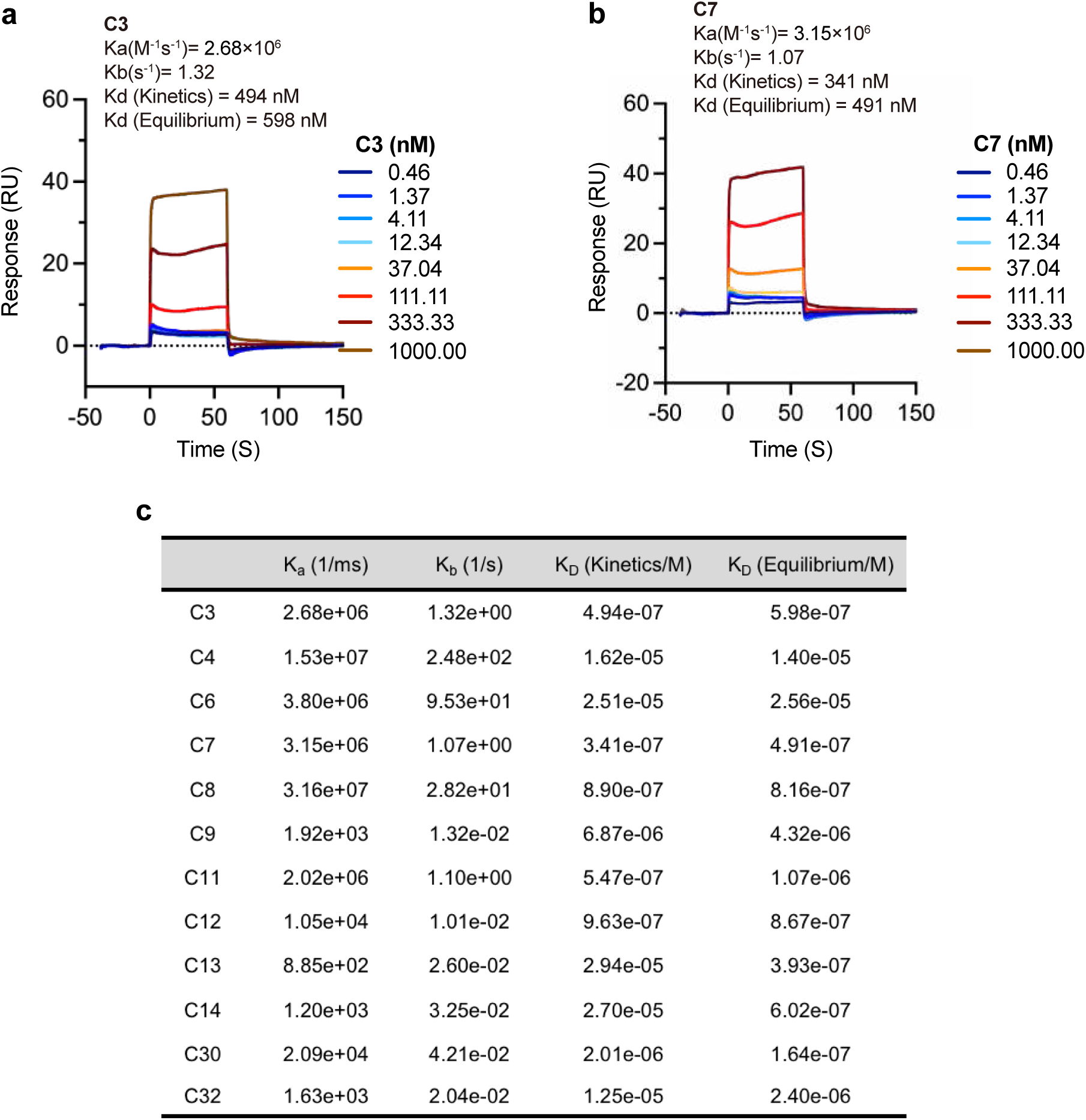
SPR binding assays for C-family compounds with immobilized NSP14. **a-b.** The binding affinity of C3 (a) and C7 (b) with immobilized NSP14, as determined by SPR. **c.** Summary of the binding affinity and kinetic parameters of C-family compounds with immobilized NSP14.

**Extended Data Figure 6.**
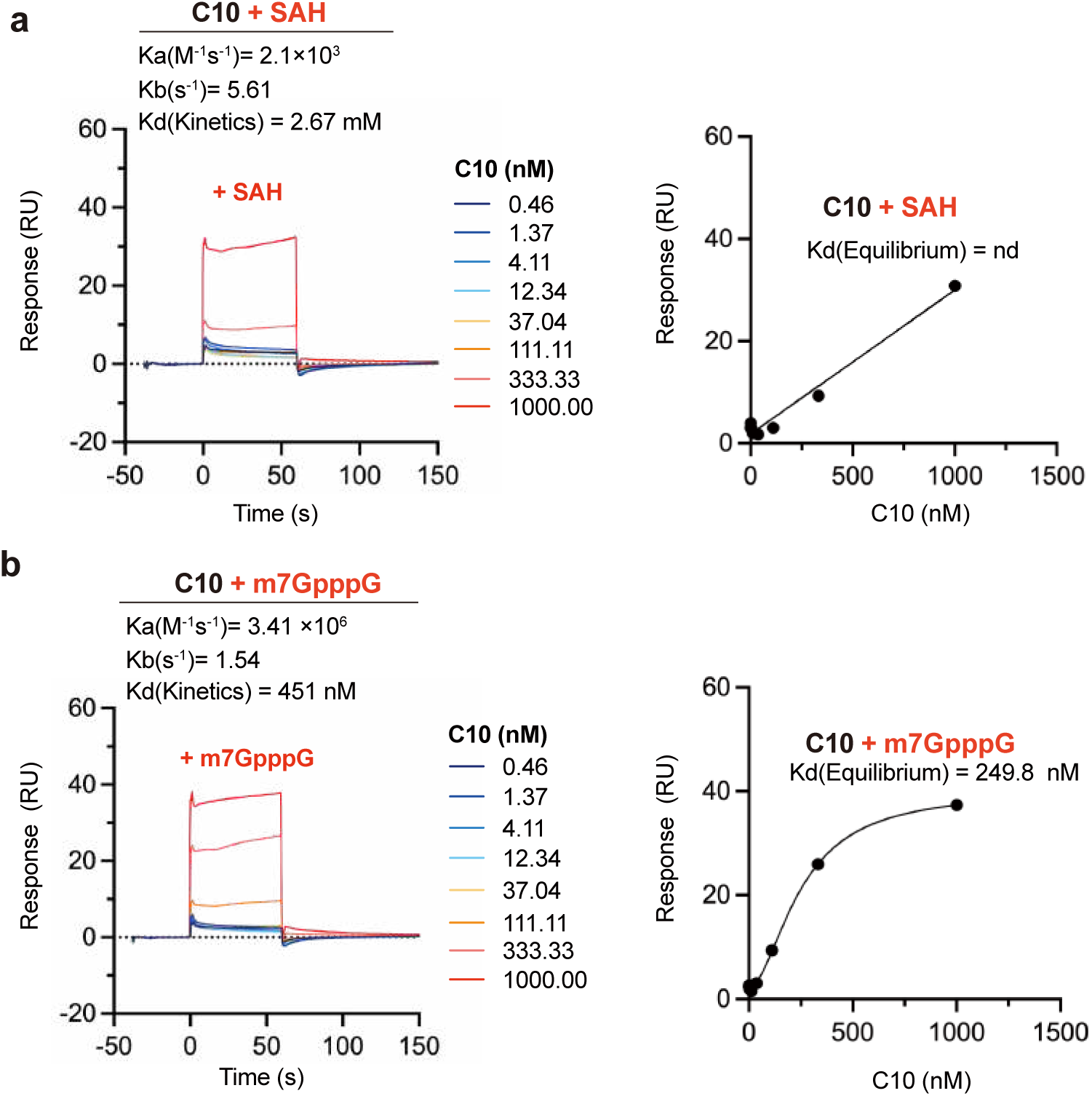
SPR binding assays for NSP14 and C10 in the presence of SAH or m7GpppG. **a-b**. SPR binding data indicating that the binding affinity of C10 with immobilized NSP14 is reduced in the presence of SAH (a) compared to m7GpppG (b), suggesting a SAH-competitive binding mode.

**Extended Data Figure 7.**
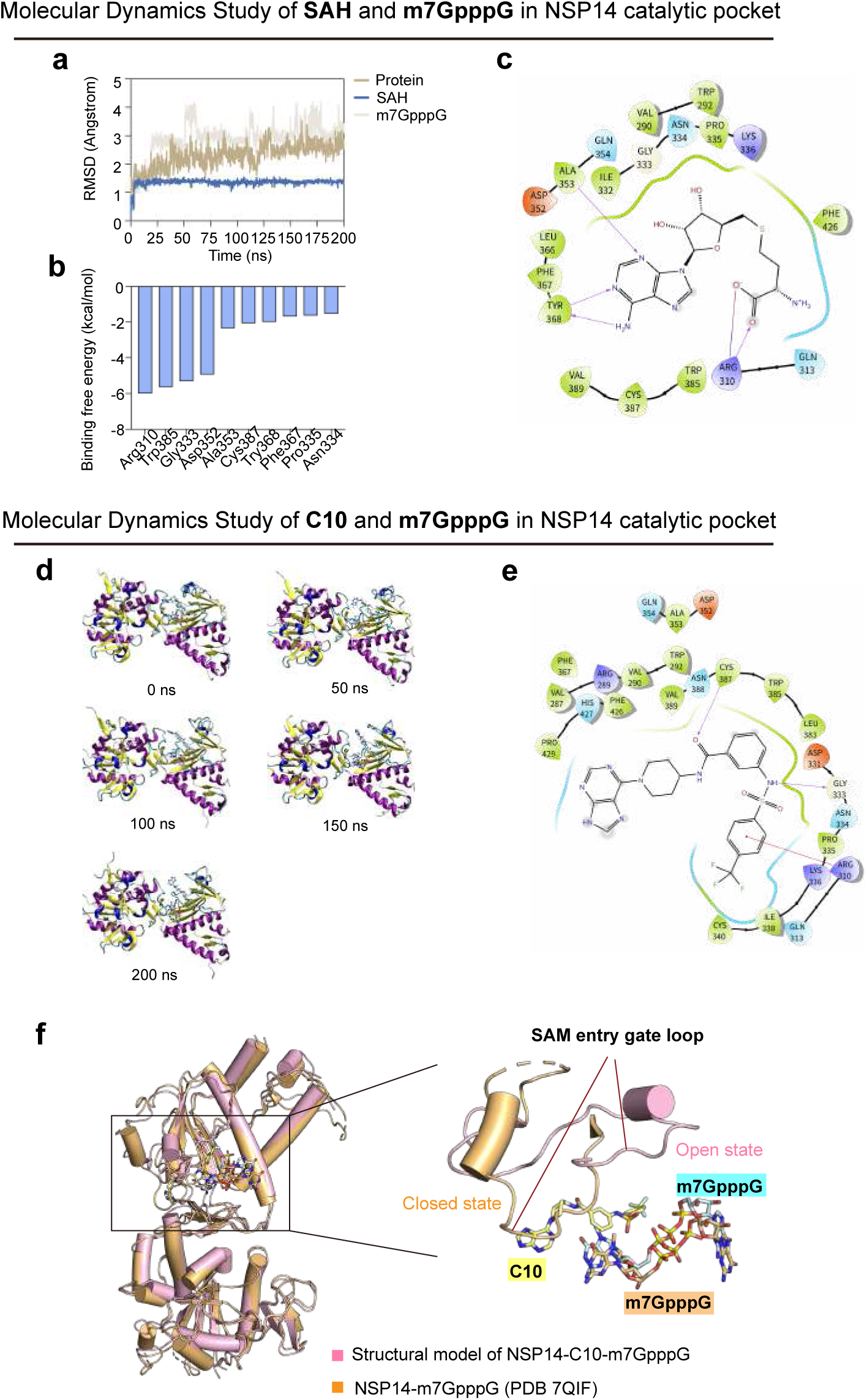
Molecular docking and MD simulations results. **a.** RMSD fluctuation of the SAH-NSP14-m7GpppG complex over the simulation time. **b.** Top 10 key residues of NSP14 responsible for binding to SAH, predicted by MM/GBSA. **c.** 2D schematic showing key residues of NSP14 involved in binding to SAH. **d.** 3D schematic of the C10-NSP14-m7GpppG complex throughout the simulation time. **e.** 2D schematic showing key residues of NSP14 involved in binding to C10. **f.** Structural alignment of the molecular docking model of NSP14-C10-m7GpppG with the crystal structure of NSP14-m7GpppG (PDB ID: 7QIF). An expanded view highlights the dynamic nature of the SAM entry gate loop, which, in its closed conformation, significantly overlaps with the core structure of C10.

**Extended Data Figure 8.**
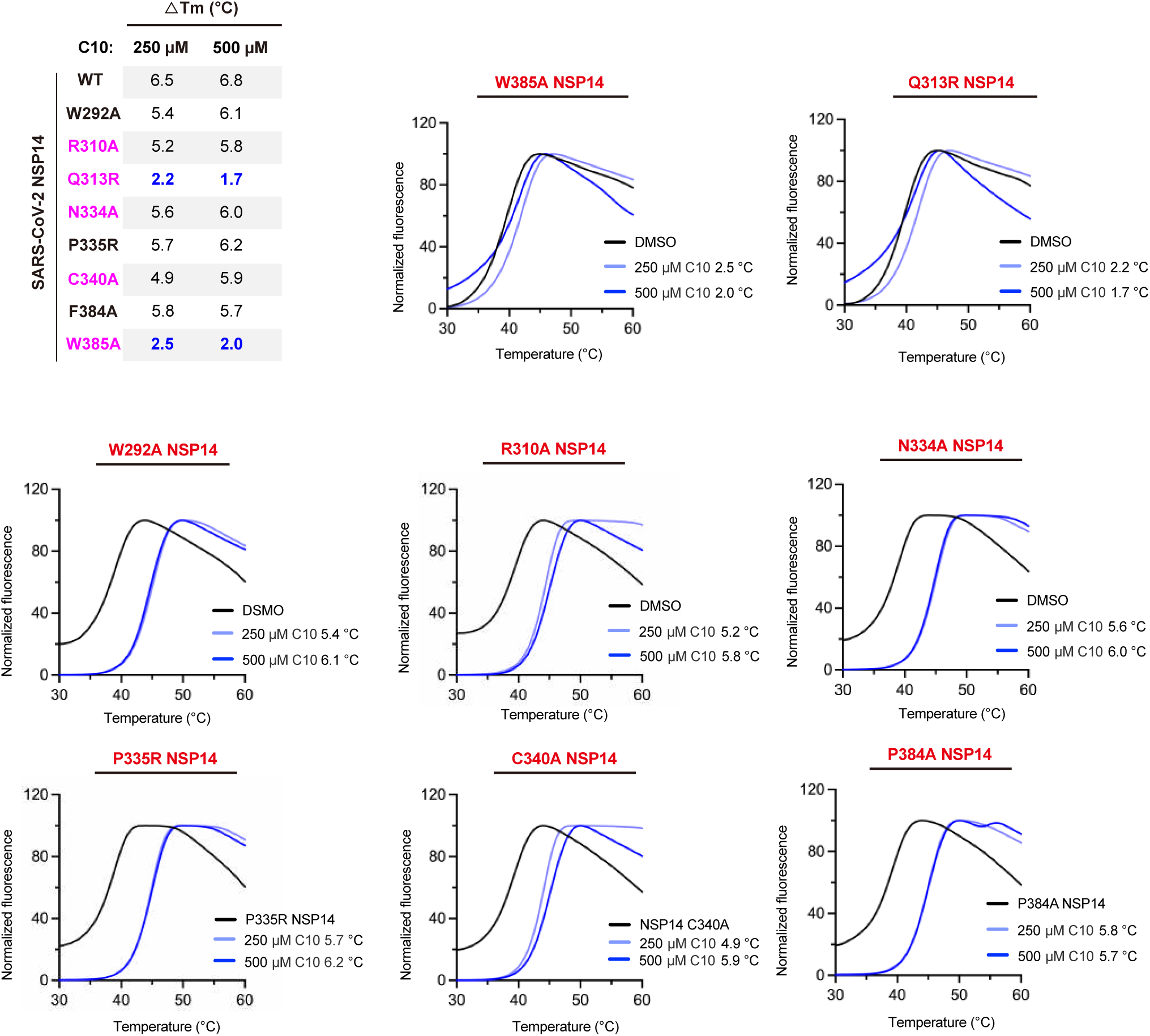
Melting temperature shifts of SARS-CoV-2 NSP14 mutants by adding 250 μM and 500 μM C10, as measured by the DSF assays.

**Extended Data Figure 9.**
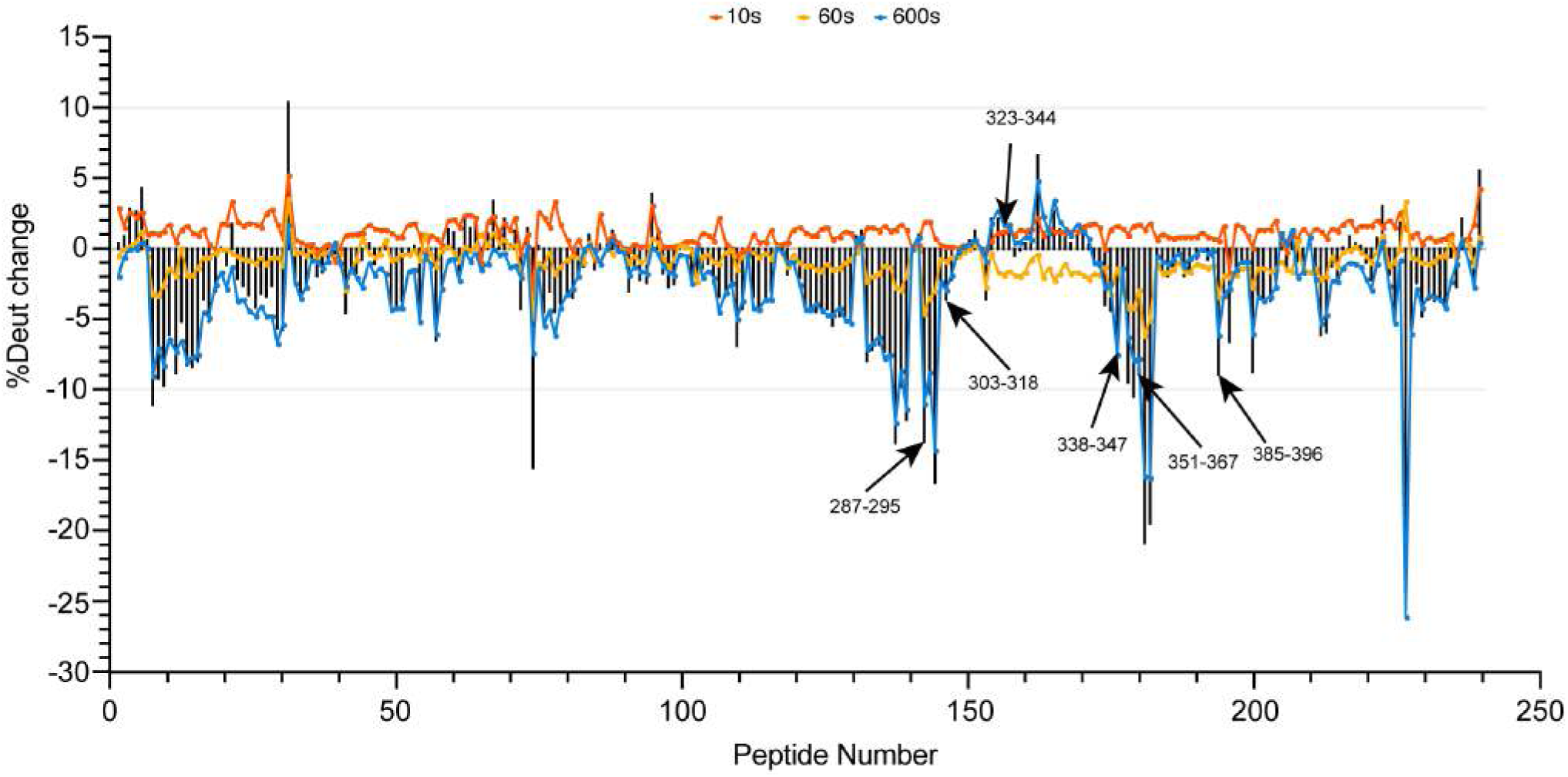
General summary of Hydrogen-Deuterium Exchange Mass Spectrometry. Line chart for the change of deuterium exchange rate is shown for all fragments after C10 binding at three time points corresponding to 10, 60 and 600 s. The bar chart represents the accumulation of the change of deuterium exchange rate for all fragments after C10 binding at 10, 60 and 600 s. %Deut change = %Deut (NSP14-C10) - %Deut (NSP14 apo).

**Extended Data Figure 10.**
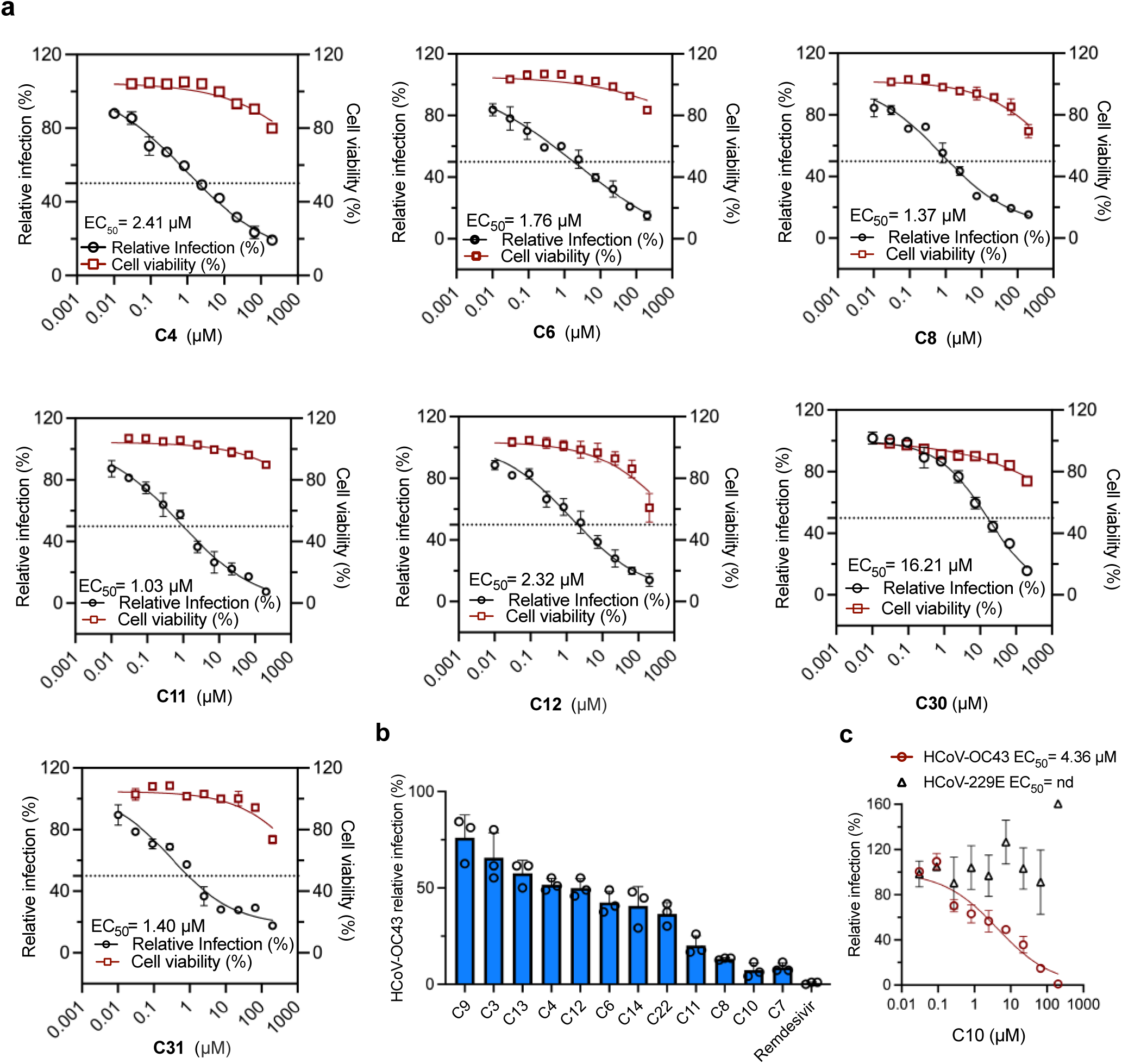
Anti-viral activities in cell-based assays of selected C-family compounds. **a.** Dose-dependent inhibition curves of selected C-family compounds against SARS-CoV-2 original strain, with the 50% cytotoxic concentration (CC_50_) values as measured in A549-ACE2 cells. Data are presented as mean ± SD from three independent replicates. **b.** Histogram showing the relative infection of HCoV-OC43 in A549-ACE2 cells treated with selected C-family compounds at 20 μM. Data are presented as mean ± SD from three independent replicates. **c.** Dose-dependent inhibition curves of C10 against HCoV-OC43 and HCoV-229E virus as measured in A549-ACE2 cells. Data are presented as mean ± SD from three independent replicates.

**Extended Data Figure 11.**
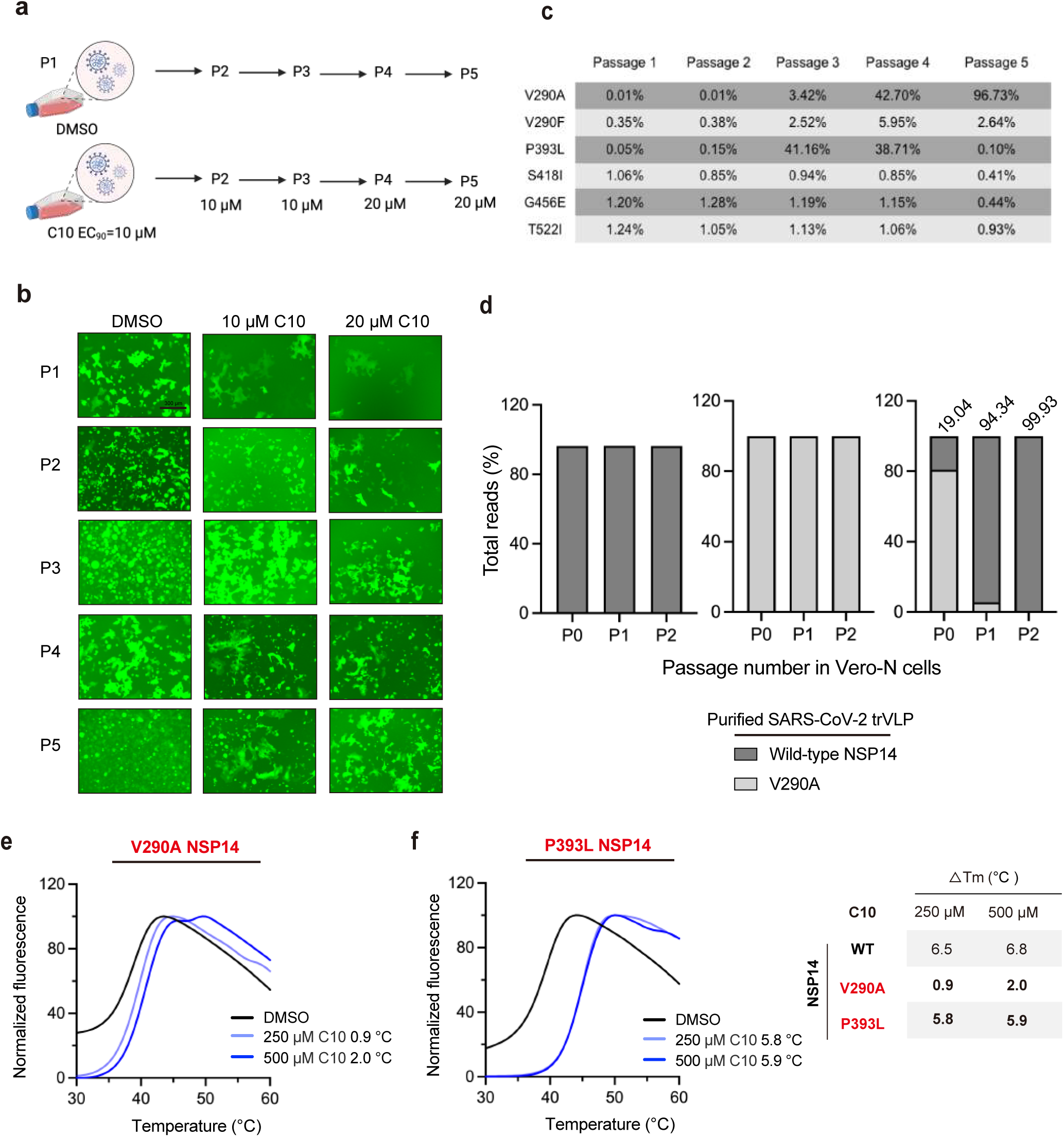
Selection of resistance mutations and fitness competition assays. **a.** Schematic of the SARS-CoV-2 trVLP passaging protocol used to select for resistance mutations to C10 in Vero-N cells. **b.** Fluorescence images showing inhibition of C10-passaged viruses after each passage by treatment with 10 μM and 20 μM C10. **c.** Table summarizing the percentage of NSP14 amino acid changes emerging in SARS-CoV-2 trVLP populations. **d.** Assessment of viral fitness of wild-type and NSP14-V290A purified SARS-CoV-2 trVLP co-cultured starting at 10:0, 0:10, or 1:9 ratio monitored over 2 passages in A549-ACE2 cells, respectively. **e-f.** Dose-dependent increase in melting temperature of NSP14 V290A and P393L mutants upon treatment with 250 μM and 500 μM C10, measured by DSF assay.

**Extended Data Figure 12.**
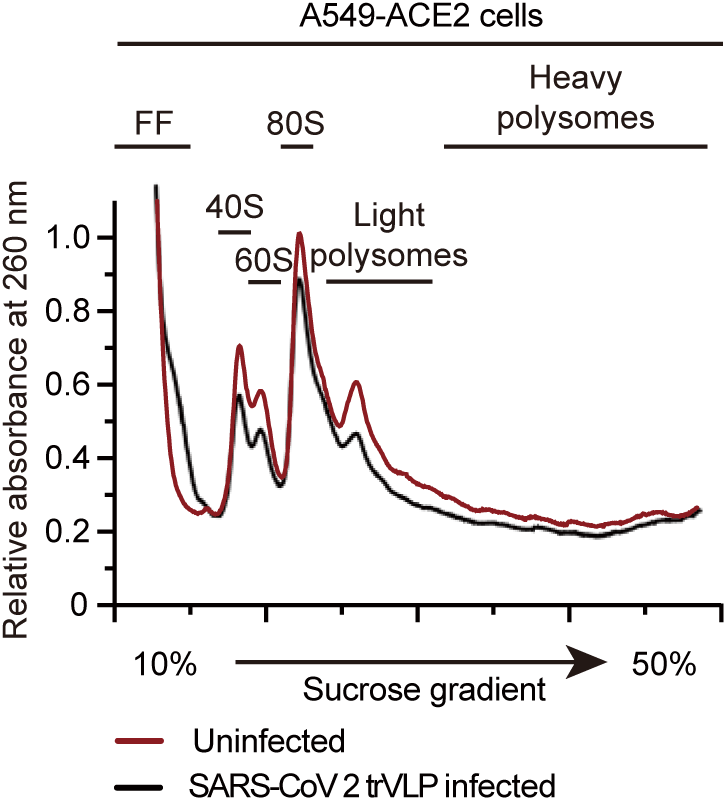
Polysome profiling of mock-infected and VLP-infected cell lysates.

**Extended Data Figure 13.**
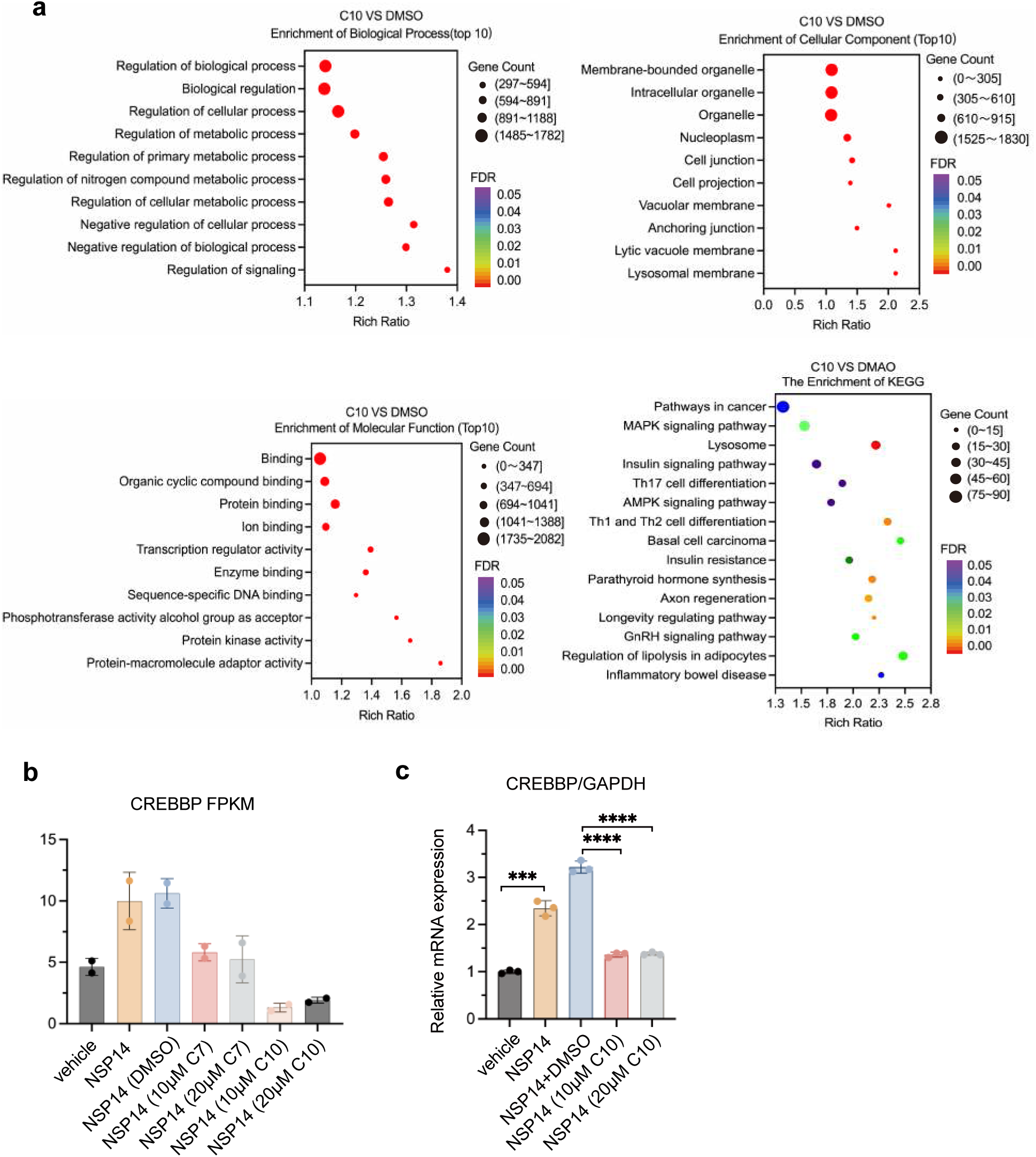
RNA-seq analyses for C10 vs DMSO in HEK293T-NSP14 cells. **a.** Bubble plots of gene signaling pathway enrichment for C10 vs DMSO in HEK293T-NSP14 cells. **b.** FPKM analysis of CREBBP for RNA-seq data from four samples: vehicle, NSP14 expression, NSP14 expression plus DMSO treatment, and NSP14 expression plus C7 (10 μM and 20 μM) treatment. Data are presented as averages of two independent experiments. **c.** Validation of CREBBP transcriptional changes by qPCR. Data are presented as mean ± SD from three replicates. Statistical significance was analyzed by unpaired t-test. P**** < 0.0001.

**Extended Data Figure 14.**
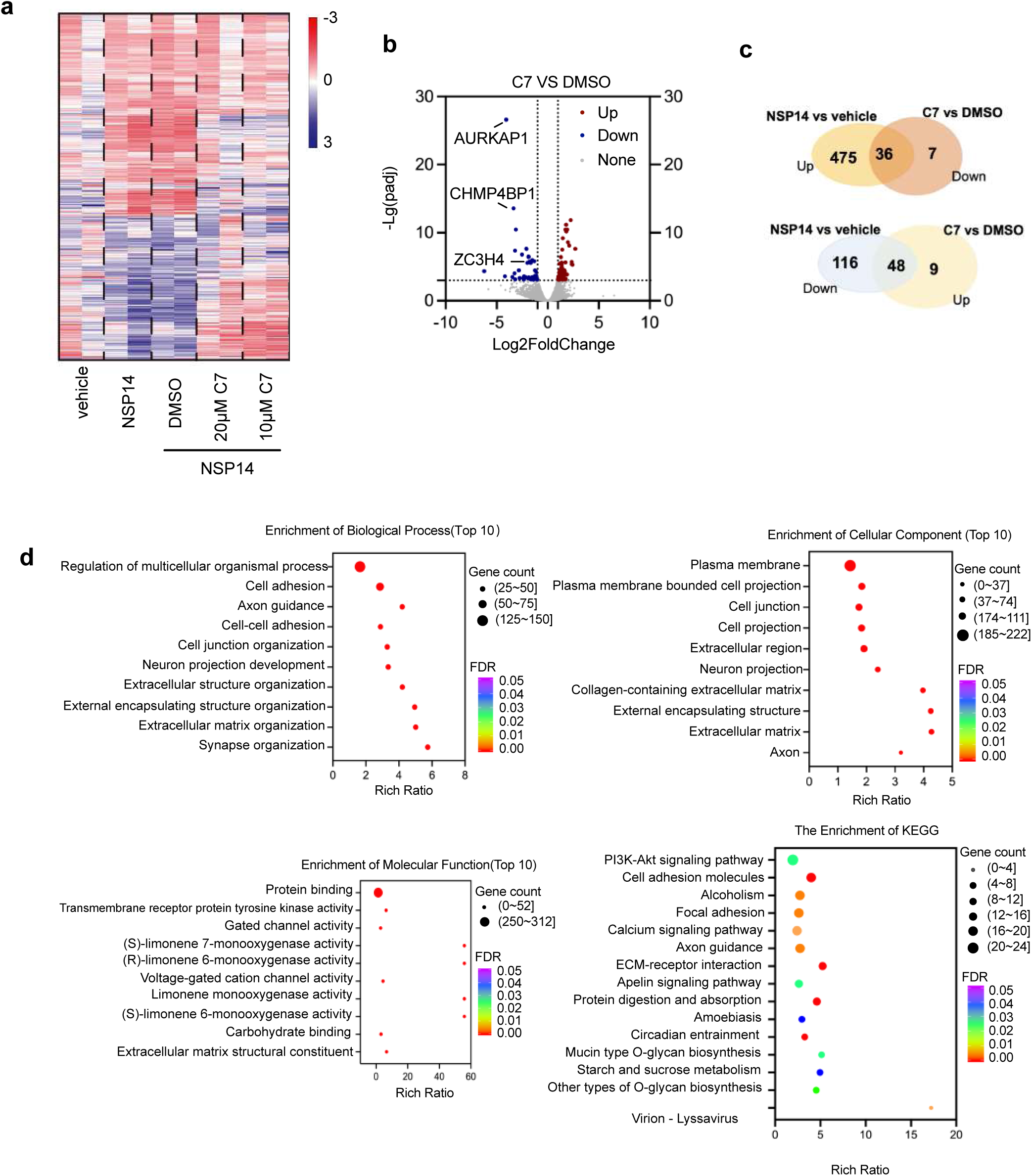
RNA-seq analyses for C7 vs DMSO in HEK293T-NSP14 cells. a. Volcano plots of RNA-seq data showing the upregulated and downregulated gene expression in HEK293-NSP14 cells treated with C7 (10 μM) vs DMSO for 48 hours. The filter condition was 2-fold change, *P*.adj < 0.001. *P*.adj refers to the adjusted P value. Data are presented as averages of two independent experiments. **b.** Cluster heatmap of differentially expressed transcription factors for RNA-seq data from four samples: vehicle, NSP14 expression, NSP14 expression plus DMSO treatment, and NSP14 expression plus C7 (10 μM) treatment. Data are presented as averages of two independent experiments. **c.** Venn diagram showing overlap of upregulated and downregulated expressed genes between NSP14 vs vehicle and 10 μM C7 vs DMSO. **d.** Bubble plots of gene signaling pathway enrichment for C7 vs DMSO in HEK293T-NSP14 cells.

**Extended Data Figure 15.**
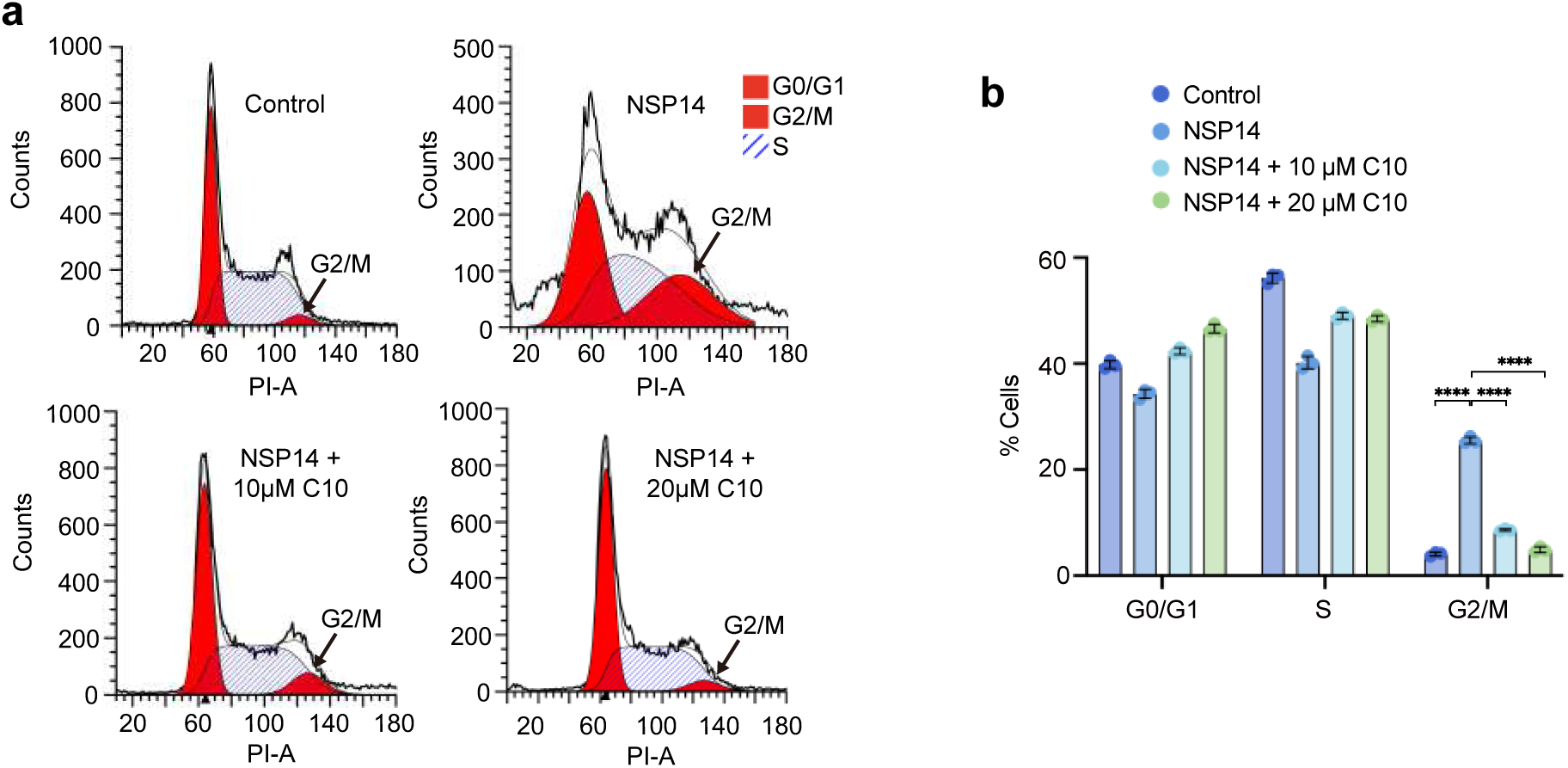
C10 recovers NSP14-triggered host cell cycle modulation. **a-b.** HEK293T cells were transfected with NSP14 and treated with C10 or not for 48 hours. After 48 hours, cells were harvested, and the cell cycle was analyzed by flow cytometry (**a**). Data from three independent experiments are shown in the column graphs (**b**). Statistical significance was analyzed by unpaired t-test. P****<0.0001.

**Extended Data Figure 16.**
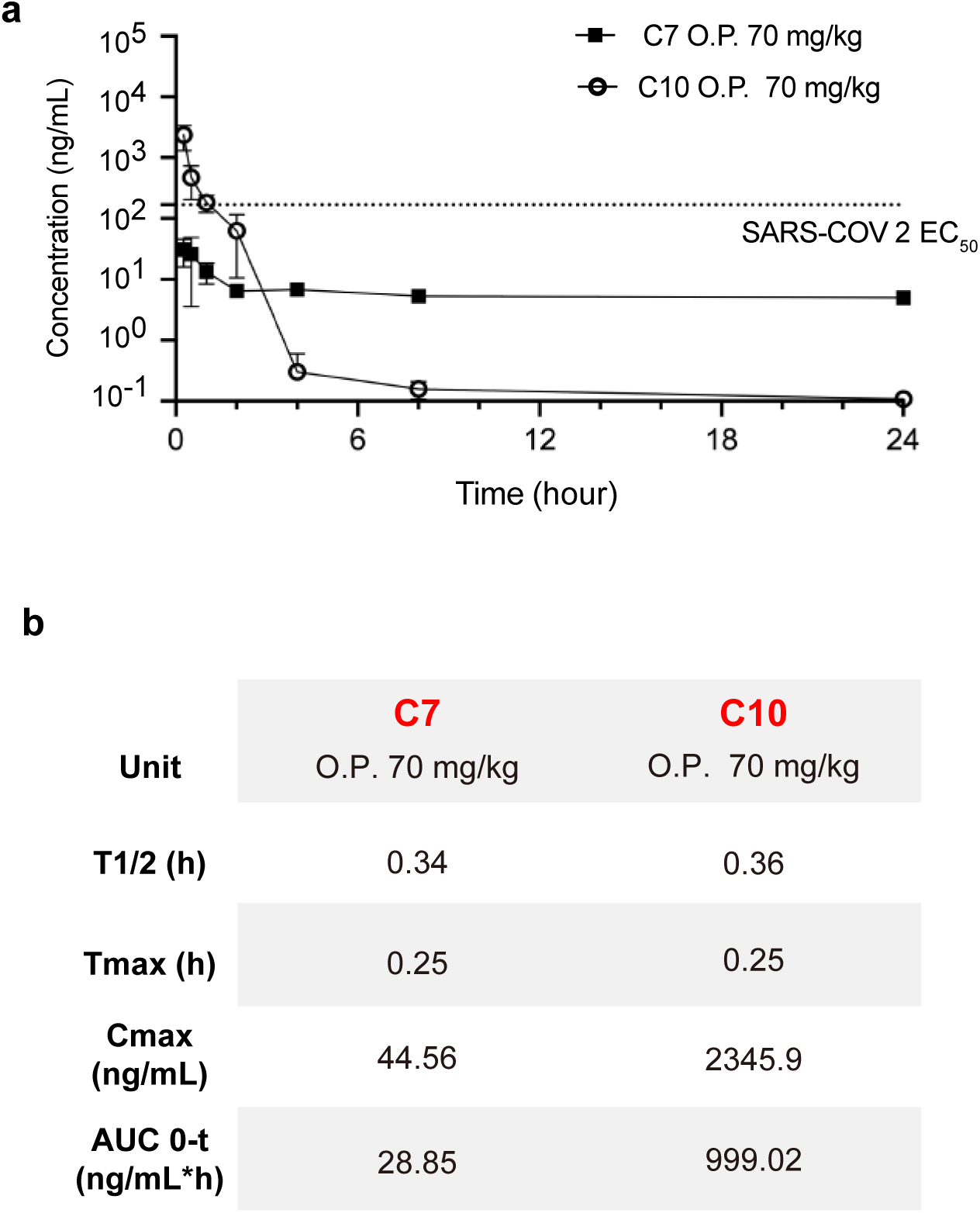
Plasma concentration-time curves for C7 and C10 in ICR mice following oral administration. **a.** Plasma drug concentration of C7 and C10 in Male ICR mice (6-8 weeks) following oral administration of 70 mg/kg of compounds in a 5% dimethyl sulfoxide and 95% cyclodextrin solution (m(cyclodextrin):m(normal saline) = 1:2) (n = 3 per group). O.P. refers to oral administration. **b.** In vitro pharmacokinetic (PK) parameters of C7 and C10.

## References

1. Li, J., Lai, S., Gao, G. F. & Shi, W. The emergence, genomic diversity and global spread of SARS-CoV-2. Nature 600, 408–418 (2021).

2. Phillips, N. The coronavirus is here to stay - here’s what that means. Nature 590, 382–384 (2021).

3. Li, G., Hilgenfeld, R., Whitley, R. & De Clercq, E. Therapeutic strategies for COVID-19: progress and lessons learned. Nat. Rev. Drug Discov. 22, 449–475 (2023).

4. Hu, Y. et al. Naturally Occurring Mutations of SARS-CoV-2 Main Protease Confer Drug Resistance to Nirmatrelvir. ACS Cent. Sci. 9, 1658–1669 (2023).

5. Iketani, S. et al. Multiple pathways for SARS-CoV-2 resistance to nirmatrelvir. Nature 613, 558–564 (2023).

6. Park, G. J. et al. The mechanism of RNA capping by SARS-CoV-2. Nature 609, 793–800 (2022).

7. Decroly, E., Ferron, F., Lescar, J. & Canard, B. Conventional and unconventional mechanisms for capping viral mRNA. Nat. Rev. Microbiol. 10, 51–65 (2011).

8. Hsu, J. C.-C., Laurent-Rolle, M., Pawlak, J. B., Wilen, C. B. & Cresswell, P. Translational shutdown and evasion of the innate immune response by SARS-CoV-2 NSP14 protein. Proc. Natl. Acad. Sci. U. S. A. 118, (2021).

9. Minkoff, J. M. & tenOever, B. Innate immune evasion strategies of SARS-CoV-2. Nat. Rev. Microbiol. 21, 178–194 (2023).

10. Pan, R. et al. N7-Methylation of the Coronavirus RNA Cap Is Required for Maximal Virulence by Preventing Innate Immune Recognition. mBio 13, e0366221 (2022).

11. Tahir, M. Coronavirus genomic nsp14-ExoN, structure, role, mechanism, and potential application as a drug target. J. Med. Virol. 93, 4258–4264 (2021).

12. Bouvet, M. et al. RNA 3’-end mismatch excision by the severe acute respiratory syndrome coronavirus nonstructural protein nsp10/nsp14 exoribonuclease complex. Proc. Natl. Acad. Sci. 109, 9372–9377 (2012).

13. Chen, Y. et al. Functional screen reveals SARS coronavirus nonstructural protein nsp14 as a novel cap N7 methyltransferase. Proc. Natl. Acad. Sci. U. S. A. 106, 3484–3489 (2009).

14. Ma, Y. et al. Structural basis and functional analysis of the SARS coronavirus nsp14-nsp10 complex. Proc. Natl. Acad. Sci. U. S. A. 112, 9436–9441 (2015).

15. Yan, L. et al. Cryo-EM structure of an extended SARS-CoV-2 replication and transcription complex reveals an intermediate state in cap synthesis. Cell 184, 184–193 (2021).

16. Yan, L. et al. Coupling of N7-methyltransferase and 3′-5′ exoribonuclease with SARS-CoV-2 polymerase reveals mechanisms for capping and proofreading. Cell 184, 3474–3485 (2021).

17. Case, J. B., Ashbrook, A. W., Dermody, T. S. & Denison, M. R. Mutagenesis of S-adenosyl-l-methionine-binding residues in coronavirus nsp14 N7-methyltransferase demonstrates differing requirements for genome translation and resistance to innate immunity. J. Virol. 90, 7248–7256 (2016).

18. Ogando, N. S. et al. The enzymatic activity of the nsp14 exoribonuclease is critical for replication of MERS-CoV and SARS-CoV-2. J. Virol. 94, 10–1128 (2020).

19. Ogando, N. S. et al. Structure-function analysis of the nsp14 N7-guanine methyltransferase reveals an essential role in Betacoronavirus replication. Proc. Natl. Acad. Sci. U. S. A. 118, (2021).

20. Canal, B. et al. Identifying SARS-CoV-2 antiviral compounds by screening for small molecule inhibitors of nsp14/nsp10 exoribonuclease. Biochem. J. 478, 2445–2464 (2021).

21. Nencka, R. et al. Coronaviral RNA-methyltransferases: function, structure and inhibition. Nucleic Acids Res. 50, 635–650 (2022).

22. Ferron, F., Decroly, E., Selisko, B. & Canard, B. The viral RNA capping machinery as a target for antiviral drugs. Antiviral Res. 96, 21–31 (2012).

23. Zaffagni, M. et al. SARS-CoV-2 Nsp14 mediates the effects of viral infection on the host cell transcriptome. eLife 11, (2022).

24. Zilecka, E. et al. Structure of SARS-CoV-2 MTase nsp14 with the inhibitor STM957 reveals inhibition mechanism that is shared with a poxviral MTase VP39. J. Struct. Biol. X 10, 100109 (2024).

25. Yan, L. et al. A mechanism for SARS-CoV-2 RNA capping and its inhibition by nucleotide analog inhibitors. Cell 185, 4347–4360.e17 (2022).

26. Yang, H. & Rao, Z. Structural biology of SARS-CoV-2 and implications for therapeutic development. Nat. Rev. Microbiol. 19, 685–700 (2021).

27. Imprachim, N., Yosaatmadja, Y. & Newman, J. A. Crystal structures and fragment screening of SARS-CoV-2 NSP14 reveal details of exoribonuclease activation and mRNA capping and provide starting points for antiviral drug development. Nucleic Acids Res. 51, 475–487 (2023).

28. Malone, B., Urakova, N., Snijder, E. J. & Campbell, E. A. Structures and functions of coronavirus replication-transcription complexes and their relevance for SARS-CoV-2 drug design. Nat. Rev. Mol. Cell Biol. 23, 21–39 (2022).

29. Kottur, J., Rechkoblit, O., Quintana-Feliciano, R., Sciaky, D. & Aggarwal, A. K. High-resolution structures of the SARS-CoV-2 N7-methyltransferase inform therapeutic development. Nat. Struct. Mol. Biol. 29, 850–853 (2022).

30. Bobileva, O. et al. Potent SARS-CoV-2 mRNA cap methyltransferase inhibitors by bioisosteric replacement of methionine in SAM cosubstrate. ACS Med. Chem. Lett. 12, 1102–1107 (2021).

31. Hausdorff, M. et al. Structure-guided optimization of adenosine mimetics as selective and potent inhibitors of coronavirus nsp14 N7-methyltransferases. Eur. J. Med. Chem. 256, 115474 (2023).

32. Stefek, M. et al. Rational design of highly potent SARS-CoV-2 nsp14 methyltransferase inhibitors. ACS Omega 8, 27410–27418 (2023).

33. Kocek, H. et al. Discovery of highly potent SARS-CoV-2 nsp14 methyltransferase inhibitors based on adenosine 5′-carboxamides. RSC Med. Chem. 15, 3469–3476 (2024).

34. Jung, E. et al. Bisubstrate Inhibitors of Severe Acute Respiratory Syndrome Coronavirus-2 Nsp14 Methyltransferase. ACS Med. Chem. Lett. 13, 1477–1484 (2022).

35. Otava, T. et al. The structure-based design of SARS-CoV-2 nsp14 methyltransferase ligands yields nanomolar inhibitors. ACS Infect. Dis. 7, 2214–2220 (2021).

36. Ahmed-Belkacem, R. et al. Potent Inhibition of SARS-CoV-2 nsp14 N 7-Methyltransferase by Sulfonamide-Based Bisubstrate Analogues. J. Med. Chem. 65, 6231–6249 (2022).

37. Devkota, K. et al. Probing the SAM binding site of SARS-CoV-2 Nsp14 in vitro using SAM competitive inhibitors guides developing selective bisubstrate inhibitors. SLAS Discov. Adv. Sci. Drug Discov. 26, 1200–1211 (2021).

38. Amador, R. et al. Facile access to 4′-(N-acylsulfonamide) modified nucleosides and evaluation of their inhibitory activity against SARS-CoV-2 RNA cap N 7-guanine-methyltransferase nsp14. Org. Biomol. Chem. 20, 7582–7586 (2022).

39. Wen, Y., et al. Identification and Evaluation of Non-Nucleosidic MTase Inhibitors against SARS-CoV-2 nsp14 with Lower-Micromolar Anti-Coronavirus Activity. ACS Infect. Dis. (2025).

40. Li, X. & Song, Y. Perspective for Drug Discovery Targeting SARS Coronavirus Methyltransferases: Function, Structure and Inhibition. J. Med. Chem. 67, 18642–18655 (2024).

41. Singh, I. et al. Structure-Based Discovery of Inhibitors of the SARS-CoV-2 Nsp14 N7-Methyltransferase. J. Med. Chem. 66, 7785–7803 (2023).

42. Basu, S. et al. Identifying SARS-CoV-2 antiviral compounds by screening for small molecule inhibitors of Nsp14 RNA cap methyltransferase. Biochem. J. 478, 2481–2497 (2021).

43. Kasprzyk, R. et al. Identification and evaluation of potential SARS-CoV-2 antiviral agents targeting mRNA cap guanine N7-Methyltransferase. Antiviral Res. 193, 105142 (2021).

44. Meyer, C. et al. Small-molecule inhibition of SARS-CoV-2 NSP14 RNA cap methyltransferase. Nature 637, 1178–1185 (2025).

45. Liao, Z. et al. DeepDock: enhancing ligand-protein interaction prediction by a combination of ligand and structure information. in 311–317 (IEEE, 2019).

46. De Wit, E., Van Doremalen, N., Falzarano, D. & Munster, V. J. SARS and MERS: recent insights into emerging coronaviruses. Nat. Rev. Microbiol. 14, 523–534 (2016).

47. Chen, Y., Wang, X., Shi, H. & Zou, P. Montelukast inhibits HCoV-OC43 infection as a viral inactivator. Viruses 14, 861 (2022).

48. Zhou, H. et al. The conformational changes of Zika virus methyltransferase upon converting SAM to SAH. Oncotarget 8, 14830 (2017).

49. Yankova, E. et al. Small-molecule inhibition of METTL3 as a strategy against myeloid leukaemia. Nature 593, 597–601 (2021).

50. San José-Enériz, E., et al. Discovery of first-in-class reversible dual small molecule inhibitors against G9a and DNMTs in hematological malignancies. Nat. Commun. 8, 15424 (2017).

51. Samrat, S. K. et al. A universal fluorescence polarization high throughput screening assay to target the SAM-binding sites of SARS-CoV-2 and other viral methyltransferases. Emerg. Microbes Infect. 12, 2204164 (2023).

52. Rona, G. et al. The NSP14/NSP10 RNA repair complex as a Pan-coronavirus therapeutic target. Cell Death Differ. 29, 285–292 (2022).

53. Zhang, J. & Zheng, Y. G. SAM/SAH analogs as versatile tools for SAM-dependent methyltransferases. ACS Chem. Biol. 11, 583–597 (2016).

54. Xiong, L. et al. Discovery of a Potent and Cell-Active Inhibitor of DNA 6mA Demethylase ALKBH1. J. Am. Chem. Soc. 146, 6992–7006 (2024).

55. Kokic, G. et al. Mechanism of SARS-CoV-2 polymerase stalling by remdesivir. Nat. Commun. 12, 279 (2021).

56. Beigel, J. H., Tomashek, K. M. & Dodd, L. E. Remdesivir for the Treatment of Covid-19 - Preliminary Report. Reply. N. Engl. J. Med. 383, 994 (2020).

57. Wang, H. et al. Epitranscriptomic m5C methylation of SARS-CoV-2 RNA regulates viral replication and the virulence of progeny viruses in the new infection. Sci. Adv. 10, eadn9519 (2024).

58. Ju, X. et al. A novel cell culture system modeling the SARS-CoV-2 life cycle. PLoS Pathog. 17, e1009439 (2021).

59. Yang, B. et al. A tissue specific-infection mouse model of SARS-CoV-2. Cell Discov. 9, 43 (2023).

60. Feng, F. et al. DAZAP2 functions as a pan-coronavirus restriction factor by inhibiting viral entry and genomic replication. bioRxiv 2025–02 (2025).

61. Becares, M. et al. Mutagenesis of coronavirus nsp14 reveals its potential role in modulation of the innate immune response. J. Virol. 90, 5399–5414 (2016).

62. Li, T.-W. et al. SARS-CoV-2 Nsp14 protein associates with IMPDH2 and activates NF-κB signaling. Front. Immunol. 13, 1007089 (2022).

63. Sui, L. et al. Host cell cycle checkpoint as antiviral target for SARS-CoV-2 revealed by integrative transcriptome and proteome analyses. Signal Transduct. Target. Ther. 8, 21 (2023).

64. Oladunni, F. S. et al. Lethality of SARS-CoV-2 infection in K18 human angiotensin-converting enzyme 2 transgenic mice. Nat. Commun. 11, 6122 (2020).

65. Dong, W. et al. The K18-human ACE2 transgenic mouse model recapitulates non-severe and severe COVID-19 in response to an infectious dose of the SARS-CoV-2 virus. J. Virol. 96, e00964–21 (2022).

